# Broad influence of mutant ataxin-3 on the proteome of the adult brain, young neurons, and axons reveals central molecular processes and biomarkers in SCA3/MJD using knock-in mouse model

**DOI:** 10.1101/2021.02.23.432266

**Authors:** Kalina Wiatr, Łukasz Marczak, Jean-Baptiste Perot, Emmanuel Brouillet, Julien Flament, Maciej Figiel

**Affiliations:** Institute of Bioorganic Chemistry, Polish Academy of Sciences, Noskowskiego 12/14, 61-704 Poznań, Poland; Université Paris-Saclay, Centre National de la Recherche Scientifique (CNRS), Commissariat à l’Energie Atomique (CEA), Direction de la Recherche Fondamentale (DRF), Institut de biologie François Jacob, Molecular Imaging Research Center (MIRCen), Neurodegenerative diseases Laboratory, 92265, Fontenay-aux-Roses, France

**Keywords:** ataxin-3, knock-in, knockin, SCA3, MJD, ataxia, spinocerebellar, CAG, polyQ, neurodegenerative, proteome, axon, vesicular transport, energy metabolism

## Abstract

Spinocerebellar ataxia type 3 (SCA3/MJD) has a polyQ etiology but, the current knowledge on molecular processes and proteins involved in pathogenesis is not sufficient to fully determine the disease mechanism and find drug targets. Proteases and deubiquitinases, such as Ataxin-3, often have a profound impact on other proteins, yet the global model picture of SCA3 disease progression on the protein level, showing the most crucial proteins and pathways in the brain, and neurons, was not investigated previously. Here, we investigated molecular SCA3 mechanism using interdisciplinary research paradigm combining SCA3 knock-in model, behavior, MRI, brain proteomics, precise axonal proteomics, neuronal energy recordings, labeling of vesicles, and inclusions and focusing in axonal compartment. We have demonstrated that altered metabolic and mitochondrial proteins in the brain and the lack of weight gain in Ki91 SCA3/MJD mice is reflected by the failure of energy metabolism recorded in neonatal SCA3 cerebellar neurons. We have determined that further, during disease progression, proteins responsible for metabolism, cytoskeletal architecture, vesicular and axonal transport proteins are disturbed, revealing axons as one of the essential cell compartments in SCA3 pathogenesis. Therefore we focus on SCA3 pathogenesis in axonal and somatodendritic compartments revealing highly increased axonal localization of protein synthesis machinery, including ribosomes, translation factors, and RNA binding proteins, while the level of proteins responsible for cellular transport, and mitochondria was decreased. We demonstrate the accumulation of axonal vesicles in neonatal SCA3 cerebellar neurons and increased phosphorylation of SMI-312 positive adult cerebellar axons, which indicate axonal dysfunction in SCA3. In summary, the SCA3 disease mechanism is based on the broad influence of mutant ataxin-3 on the neuronal proteome. Processes central in our SCA3 model include disturbed localization of proteins between axonal and somatodendritic compartment, early neuronal energy deficit, altered neuronal cytoskeletal structure, an overabundance of protein synthetic machinery in axons.

## Introduction

Spinocerebellar ataxia type 3, also known as Machado-Joseph disease (SCA3/MJD), is a neurodegenerative, late-onset, genetic disorder caused by the expansion of CAG repeats in the coding region of the ATXN3 gene (>50 in patients) (McLoughlin et al., 2020). This mutation leads to an expanded polyQ tract in the ataxin-3 protein, causing a toxic gain of function (Orr and Zoghbi, 2007). SCA3 patients most typically display imbalance, motor incoordination, and neurodegeneration in the cerebellum, brainstem, spinal cord, and cerebral cortex (Bettencourt and Lima, 2011). Although the disease-causative mutation is known, the exact molecular and cellular mechanisms of SCA3 remain unclear. Several lines of evidence point to the disruption of axon organization and, therefore, connections between brain structures as one of the main traits in SCA3 pathogenesis (de Rezende et al., 2015; Farrar et al., 2016; Lu et al., 2017). Moreover, axonal impairment was observed in CNS and in peripheries in the form of white mater defects on MRI and peripheral axonal neuropathy in SCA3 patients (D’Abreu et al., 2009; Graves and Guiloff, 2011; Rezende et al., 2018). Axonal impairment may be related to the fact that mutant Ataxin-3 aberrantly interacts with microtubules affecting cytoskeleton dynamics in neurons and forms inclusions in axons (Mazzucchelli et al., 2009; Seidel et al., 2010; Chen et al., 2012). Also, mitochondrial impairment is a component of axonal damage in neurodegeneration, and likely plays a role in SCA3 pathogenesis (Da Silva et al., 2019). We recently demonstrated that gross transcriptional changes are absent in the pre-symptomatic SCA3 brain of the Ki91 mouse model; however, dysregulations of proteins and phosphoproteins occurred in these animals, pointing at protein homeostasis as a more general and early trait in disease pathogenesis (Switonski et al., 2015; Wiatr et al., 2019).

The ataxin-3 protein is a particular type of protease, a deubiquitinase, and it has been previously demonstrated that the function in the mutant version of the protein is compromised (Neves-Carvalho et al., 2015), which should have a significant influence on the brain proteome. Therefore, the investigation of proteins seems to be the strongest candidate to discover pathogenic mechanisms or identify the cluster of disease mechanisms. The number of known biomarkers and proteins reliably linked to SCA3 pathogenic processes is still relatively low. Surprisingly, the global model picture of SCA3 disease progression on the protein level, showing the most crucial proteins and pathways in the brain, and neurons, has not been generated and published previously. The missing knowledge about the proteome may be one reason for the incomplete understanding of the disease mechanism and lack of cure. Therefore in the present work, we aimed to identify protein dysregulations and pathogenic processes in SCA3 on various levels, *in vitro,* and *in vivo*. We designed a research pipeline consisting of three complementary proteomic approaches, mouse behavior, and necessary functional validation assays to unveil with increased resolution the most critical processes influencing the SCA3 pathogenesis. The pipeline combines proteomics paralleled to behavioral milestones of SCA3 Ki91 and correlative proteomics for displaying the proteome linked to disease severity. In the third and the most focused approach, we used targeted proteomics of somatodendritic and axonal compartments from young SCA3 neurons. Our results indicate that the SCA3 disease mechanism is based on the broad influence of mutant ataxin-3 on the neuronal proteome affecting classes of proteins responsible for several central molecular processes. These include disturbed localization of proteins between axonal and somatodendritic compartment, early neuronal energy deficit, altered neuronal cytoskeletal structure, an overabundance of protein synthetic machinery in axons, and altered vesicular trafficking pointing at affected axonal maintenance.

## Results

### 1. The SCA3 in Ki91 mice gradually progresses and demonstrates discrete early and late phenotypes

We have previously demonstrated that by 2 months of age, Ki91 mice show no motor symptoms or other behavioral abnormalities, and we additionally defined the spectrum of Ki91 motor deficits using 14-month-old animals (Wiatr et al., 2019). Upon completing our longitudinal behavioral tests (4-18 months), we now demonstrate that the homozygous Ki91 model gradually develops disease symptoms resembling disease progression in SCA3 patients. Based on the large volume of longitudinal behavioral data (Fig. 1), we have distinguished three stages of behavioral symptom development in Ki91 animals (n = 36). The first stage consists of progressive failure to gain body weight (4 month-old) followed by the early symptomatic stage (12-month-old), characterized by gait ataxia and motor incoordination. The final, symptomatic phase (18-month-old animals) is characterized by severe loss of balance and coordination, gait ataxia, dystonia, and muscle weakness (Fig. 1A). Reduced rate of body weight gain was observed as a first symptom in 4-month-old Ki91 mice (P<0.05; two-sample t-test), and the difference between Ki91 and wildtype littermates increased with age (P<0.001; two-sample t-test, two-way ANOVA, Bonferroni; Fig. 1B). At the age of 8 months, Ki91 mice stopped gaining weight, and their weight was stable across further ages (Fig. 1B) while the WT animals still regularly gained weight. First motor symptoms occurred in 12-month-old animals, which performed worse in the elevated beam walk (Fig. 1C). Ki91 took more time to traverse rods (diameter: 35, 28, 21, and 17 mm) and committed more foot slips while performing the task (P<0.05; two-way ANOVA, Bonferroni). As the disease progressed, 14-month-old Ki91 demonstrated overall deterioration of phenotype, including gait disturbances, loss of coordination, and balance, and several Ki91 animals demonstrated hindlimb clasping and kyphosis in the scoring test (P<0.0001; one- and two-way ANOVA, Bonferroni) (Fig. 1E). Next, 16-month-old Ki91 mice needed more time to turn on a rod (diameter: 28, 17, 10, and 9mm) in the elevated beam walk test (P<0.05; two-way ANOVA,

**Figure 1.**
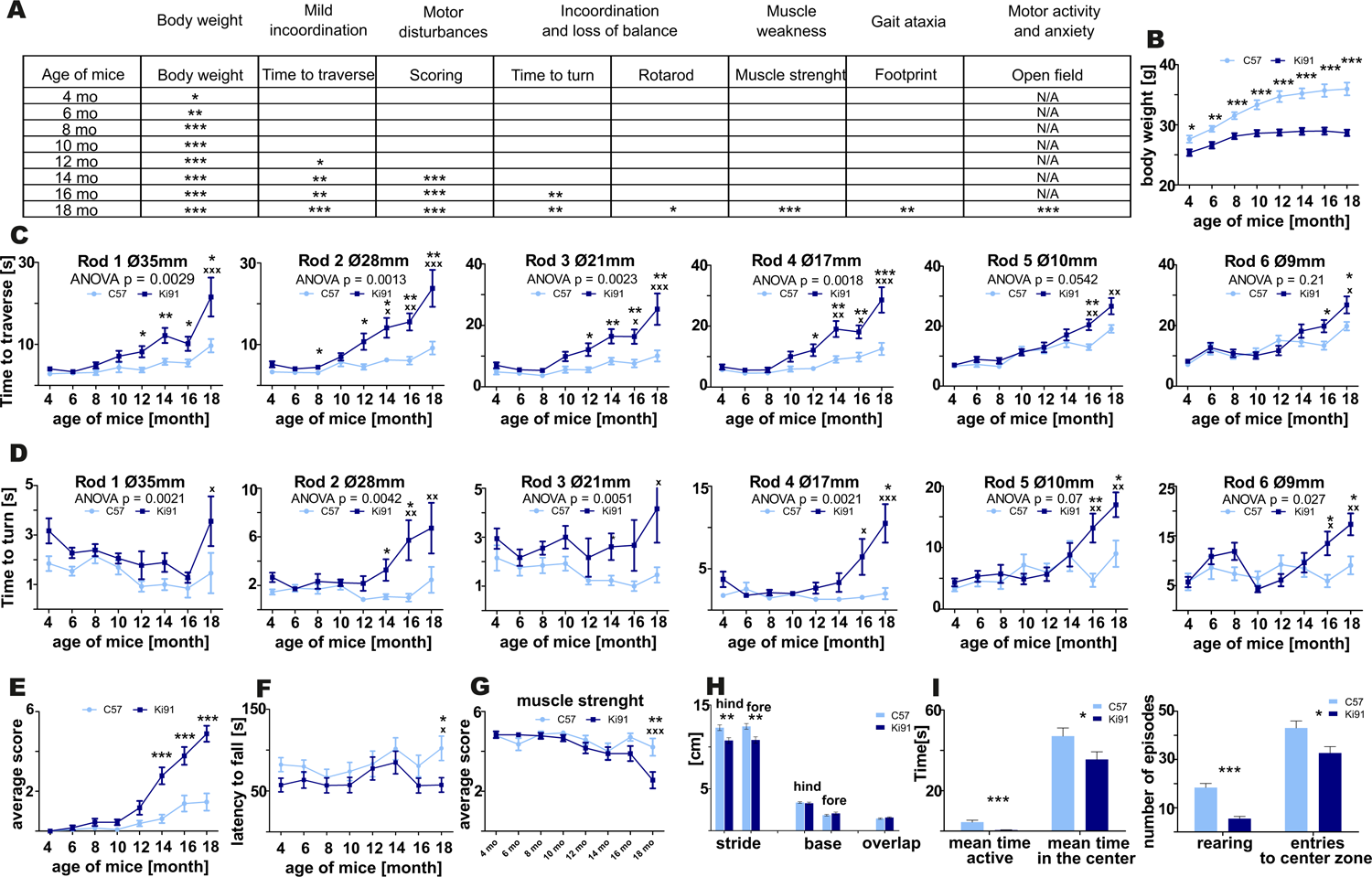
Progressive motor deficits and other disease symptoms in Ki91 SCA3/MJD mice. Ki91 mice presented a gradual decline in several motor and non-motor functions measured by elevated beam walk, rotarod, monitoring of body weight, muscle weakness, and other tests. (A). Progressive reduction of body weight gain was observed starting from 4-month-old mice; on average, mut/mut animals at 12 months of age displayed 16.4 % less body weight compared to WT animals (p<0.001; two-sample t-test) (B). In the elevated beam walk test (C-D) “time to turn” and “traverse time” parameters were measured on 6 rods with decreasing diameter (diameter of rods are indicated by Ø in mm). 12-month-old Ki91 mice needed significantly more time to traverse on rods 1-4, whereas older 16-month-old mice also needed more time on rods 5-6. (C). 16-month-old animals needed more time to turn on all rods (D). In the scoring test, 14-month-old Ki91 mice presented symptoms characteristic for SCA3: incoordination, gait disturbances, kyphosis, and hind limb clasping (E). 18-month-old mice showed motor incoordination in accelerated rotarod (4 to 40 rpm in 9.5 min) (F), muscle weakness (G), and differences in stride length in footprint test (H). 18-month-old mice also presented a cognitive deterioration marked by a decreased amount of time spent in the center zone in the Open field test and a decreased number of rearing (I). 2-way ANOVA with Bonferroni post-hoc test (p ≤ 0.05; total number of biological replicates: n=36, n=18 per genotype), error bars: SEM. Asterisks denotes a 2-way ANOVA calculated separately for each test consisted of 4 days at each age (*P < 0.05, **P < 0.01, ***P < 0.001); x-symbol represent 2-way ANOVA calculated after completion of testing all ages (^x^P < 0.05, ^xx^P < 0.01, ^xxx^P < 0.001).

Bonferroni), showing loss of balance (Fig. 1D). Besides, 18-month-old Ki91 mice demonstrated incoordination in the Rotarod (P<0.05; two-way ANOVA, Bonferroni; Fig. 1F), reduced muscle strength (P<0.001; one- and two-way ANOVA, Bonferroni; Fig. 1G), shorter stride, which indicated further gait disturbances in the footprint test (P<0.001; two-sample t-test, ANOVA, Bonferroni; Fig. 1H), and decreased activity and anxiety in the open field test (P<0.05; two-sample t-test; Fig. 1I). In this last test, Ki91 mice showed significantly less time spent moving (P<0.0001; two-sample t-test), as well as decreased vertical activity (rearing) (P<0.0001; two-two-sample t-test), a smaller number of the entries, and less time spent in the center zone (P<0.05; two-sample t-test). Since mice showed differences in body weight, we ensured that no correlation existed between body weight and behavioral tests in each age to exclude the effect of the animals’ weight on the results (P<0.05, correlation test).

### 2. Ki91 SCA3/MJD mice demonstrate multiple brain region atrophy and the presence of Atxn3 inclusions throughout the brain

MRI was used on ex vivo brains of 18 months old animals (n = 6 per genotype) to measure brain volumes in Ki91 and WT mice (Fig. 2A,B). As measured by MRI image segmentation, whole-brain volume showed significant global atrophy of the brain in Ki91 mice (−7%, p<0.001; Fig. 2A). When looking at regions volume, several structures seem particularly atrophied as parietal-temporal cortex (−11%, p<0.01), entorhinal, piriform and motor cortexes (−11% each, p<0.05), corpus callosum (−7,5%, p<0.05), striatum (−11%, p<0.05), septum (−11%, p<0.01), pons (−12%, p <0.01) and hypothalamus (−8%, p<0.05) (Fig. 2B). There was no significant atrophy in the hippocampus, and cerebellar atrophy was not determined due to MRI coil performance at the end of the brain. Fractional Anisotropy (FA) was also investigated as a biomarker of the integrity of tissue organization. FA was significantly decreased in dentate gyrus (−17%, p<0.01), and stratum granulosum (−19%, p<0.05) (Fig. 2C).

**Figure 2.**
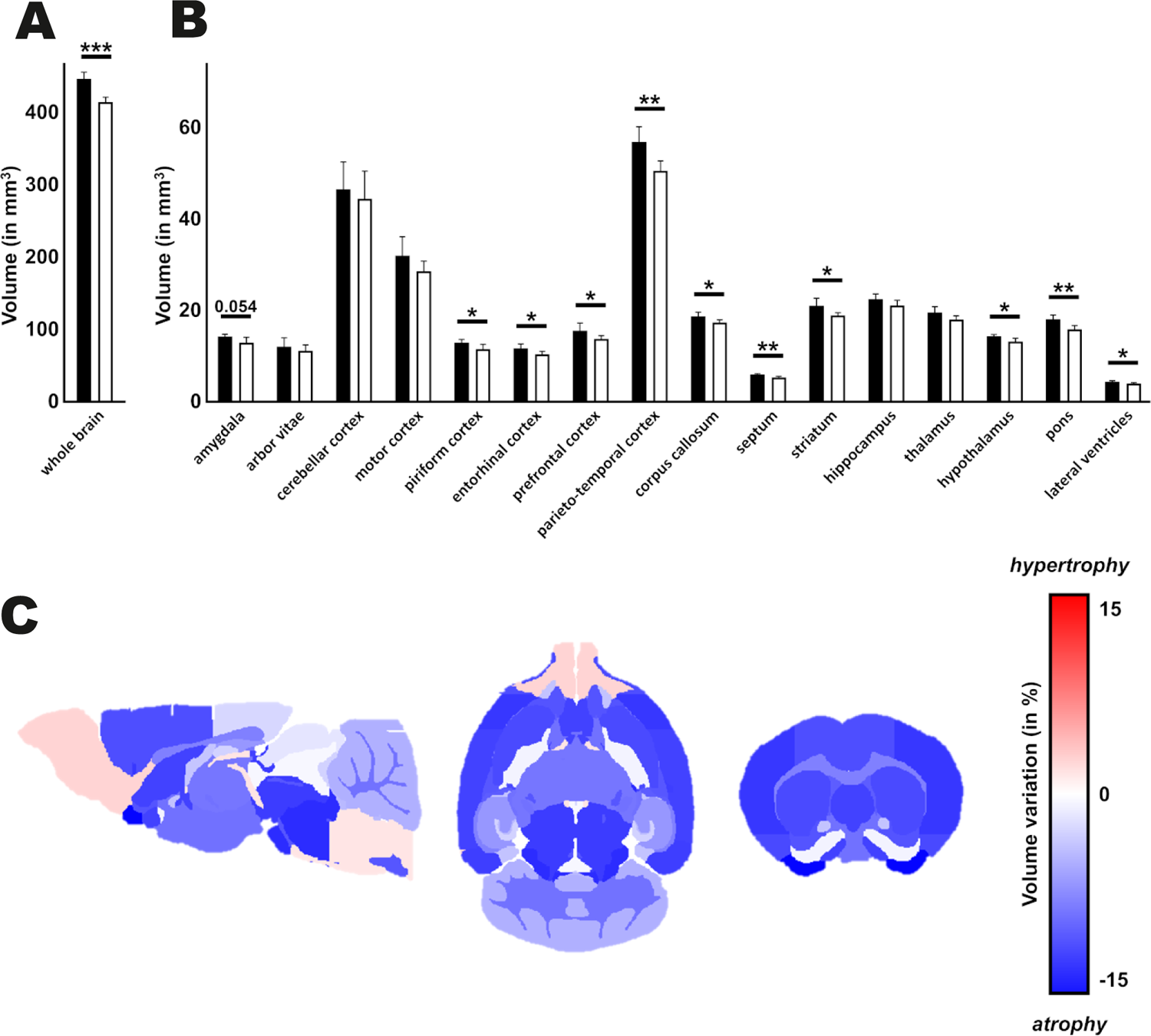
Atrophy of multiple regions including white matter in the Ki91 SCA3/MJD brain. MRI image segmentation was used to measure whole brain volume *ex vivo* of 18-month-old Ki91 and WT mice (n = 6 per genotype) and revealed significant global atrophy (−7%, p<0.001) (A) and reduction of volume of many regions, such as parieto-temporal cortex (−11%, p<0.01), entorhinal, piriform and motor cortexes (−11% each, p<0.05), corpus callosum (−7,5%, p<0.05), striatum (−11%, p<0.05), septum (−11%, p<0.01), pons (−12%, p <0.01) and hypothalamus (−8%, p<0.05) (B). Atrophy of many brain regions might suggest impaired connections between those structures. The gradient of red color is for hypertrophy, the gradient of blue color for the atrophy. Two-sample t-test (*P < 0.05, **P < 0.01, ***P < 0.001), error bars: SEM.

Next, we have performed brain sections from 18-month-old Ki91 and staining using the anti-ataxin-3 antibody (Fig. 3). We found that all brain areas, including the cerebral cortex, striatum, midbrain, hippocampus, cerebellum, DCN, and pons, demonstrated intensive positive signals in the form of both inclusions and diffuse staining of cellular structures (n=4; Fig. 3 A-D). In particular, intense anti-ataxin-3 staining was present in cell nuclei, and many cells demonstrated large inclusions (Fig. 3A). Interestingly, a significant number of smaller and larger ataxin-3-positive inclusions were scattered along the neurites (particularly axons labeled by SMI-312 axonal marker) in the cerebellum (Fig. 3 C,D). Although we could not measure cerebellum atrophy by MRI, we determined that the inclusions are rich in the cerebellum, indicating an intense pathogenic process. In the cerebellum, the large inclusions were predominantly present in the DCN area, white matter, and in some cells located in the granular layer (Fig. 3B). Considering widespread brain atrophy and the inclusions in all regions, we reasoned that the brain pathogenic SCA3 processes are multi-region and multi-thread. Therefore, to identify molecular SCA3 pathogenesis by our 3-way proteomic approach, we selected two brain regions: the cerebellum and cerebellar cortex. Such a selection of samples involves both classically reported SCA3 pathogenesis in the hindbrain and SCA3 pathogenesis in more frontal parts of the brain.

**Figure 3.**
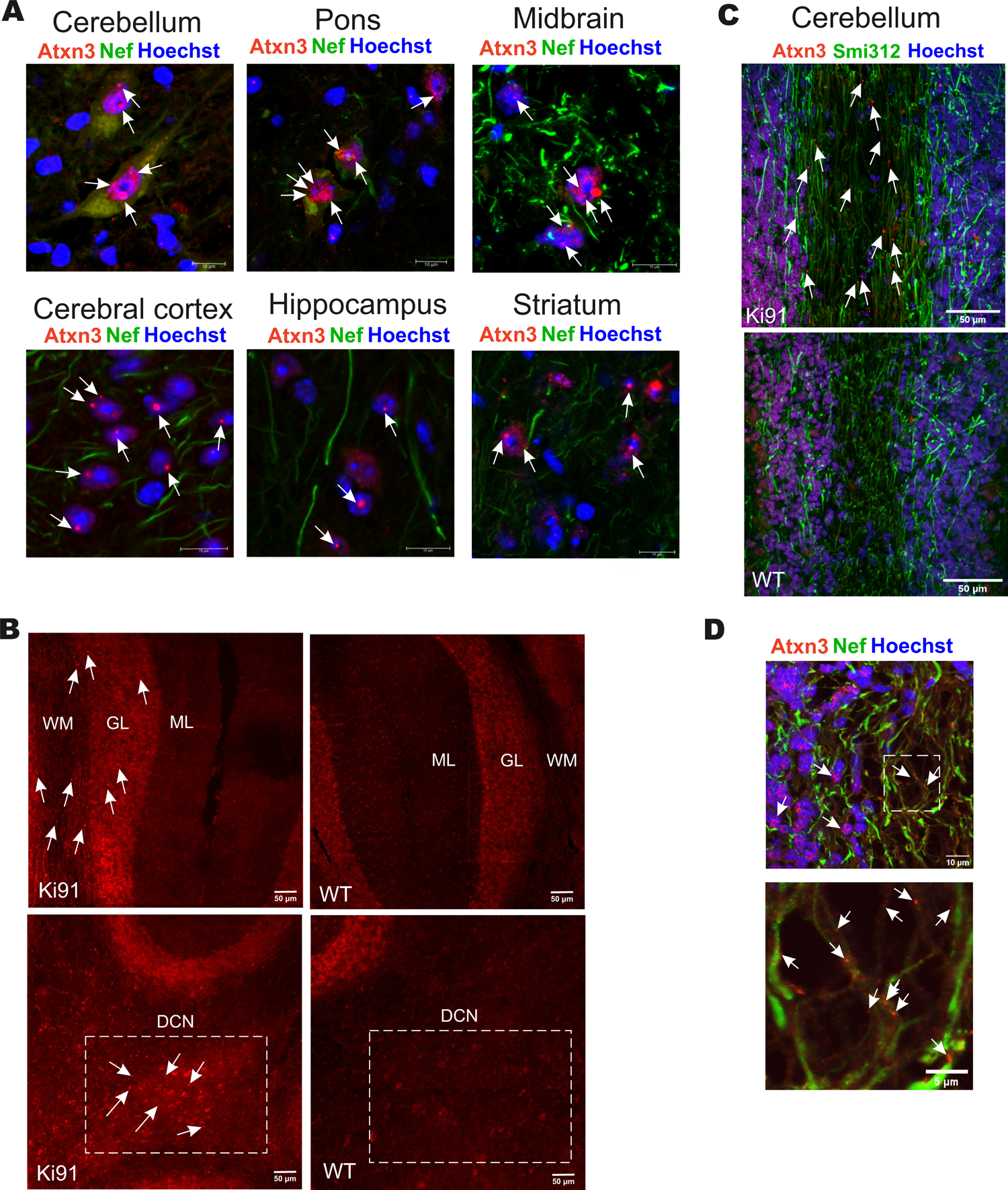
The presence of multi-region intra-nuclear and intra-axonal inclusions in Ki91 SCA3/MJD brain. The ataxin-3 immunostaining of the 18-month-old Ki91 brain sections revealed a large number of cells within the cerebellum, midbrain, pons, cerebral cortex, hippocampus, and striatum with ataxin-3 (red; rabbit anti-ataxin-3 antibody) localizing mainly in the cell nucleus (blue; Hoechst 33342) (A). White arrows indicate large intra-nuclear inclusions; however, many smaller inclusions throughout the brain were present. A rich representation of Atxn3 aggregates (red; rabbit anti-ataxin-3 antibody) was present in DCN, granular layer (gl), and white matter (wm) of the Ki91 cerebellum (B, C). A significant number of small and large ataxin-3 inclusions were detected in the neurites (green; Smi-32, pan-neuronal or Smi-312, axon-specific antibody) of the Ki91 cerebellum (C, D). White box and arrows in (D) mark the magnified area containing multiple intra-axonal inclusions of Atxn3 (lower panel). Scale bars: 50 μm (B, C); 10 μm (A); 5 μm on inserts (D). N=4 biological replicates; at least 4 pictures per brain region of each kind were collected.

### 3. Proteomics parallel to behavioral milestones implicates disturbed metabolism, cytoskeleton, and vesicular trafficking in Ki91 SCA3/MJD mice

Parallel to behavioral experiments and based on the occurrence of symptoms in behavioral tests, we have selected four mouse ages (4, 10, 12, 14-month-old, n = 4 per genotype) to analyze proteomic changes in the cerebral cortex (entire region collected for analysis) and cerebellum. In detail, 4-month-old Ki91 animals displayed reduced body weight gain; a 10-months stage was just before the onset of motor symptoms, while 12 and 14-month-old mice showed a decline in motor functions (Fig.1); therefore, the protein dysregulation in the ages would be most relevant. The proteomic analyses in the cerebellum and the cerebral cortex identified between 1265 and 2645 unique proteins, respectively. The PCA mostly demonstrated distinct clustering of the Ki91 and wildtype datasets (Suppl. Fig. 1A, B). Altogether, we identified 115, 126, 212, and 75 proteins that were significantly dysregulated (p<0.05; two-sample t-test) in the cerebellum of 4-, 10-, 12- and 14-month-old Ki91 mice, respectively (Suppl. Table 1, Suppl. Fig. 1C). In comparison, 178, 89, 170, and 279 proteins were significantly dysregulated (p<0.05; two-sample t-test) in the cerebral cortex of 4-, 10-, 12- and 14-month-old Ki91 animals, respectively (Suppl. Table 2, Suppl. Fig. 1C). Of note, the most overlap in dysregulated proteins was between 12- and 14-month-old Ki91 mice in the cerebral cortex (53 proteins) and 10- and 12-month-old animals in the cerebellum (20 proteins). Since the age between 10 and 14 in SCA3 Ki91 mice represent the phase during which motor symptoms are developed (Fig. 1), the observed protein dysregulation at these ages may yield potentially essential biomarkers of the advanced disease stage.

In order to reveal molecular pathways and cellular compartments, which might contribute to the SCA3 pathogenesis, we performed an analysis of dysregulated proteins using the ConsensusPath database (CPDB; pathways and GO Cellular Compartment (CC) for each age; level 4 and 5; q <0.01). The overrepresentation analysis in CPDB demonstrated that biological pathways that contained the highest number of dysregulated proteins (p<0.05; two-sample t-test) in all tested brain regions and ages of Ki91 were mostly related to energy metabolism, synaptic transmission, vesicular transport, regulation of cytoskeletal functions and protein metabolism (Fig. 4A-D). Coherently, the primary subcellular localization of dysregulated proteins from all tested brain regions and ages of Ki91 consisted of vesicles, axons, cytoskeleton, and mitochondria (Fig. 4A, B). For simplicity, we have arbitrarily grouped pathways and cellular compartments identified in CPDB into five categories for the cerebral cortex (Fig. 4A) and cerebellum (Fig. 4B) based on their biological relations and similarity. Lists of proteins with the highest log2-fold changes (log2-FC) belonging to 5 categories are presented for the cerebral cortex (Fig. 4C) and cerebellum (Fig. 4D).

**Figure 4.**
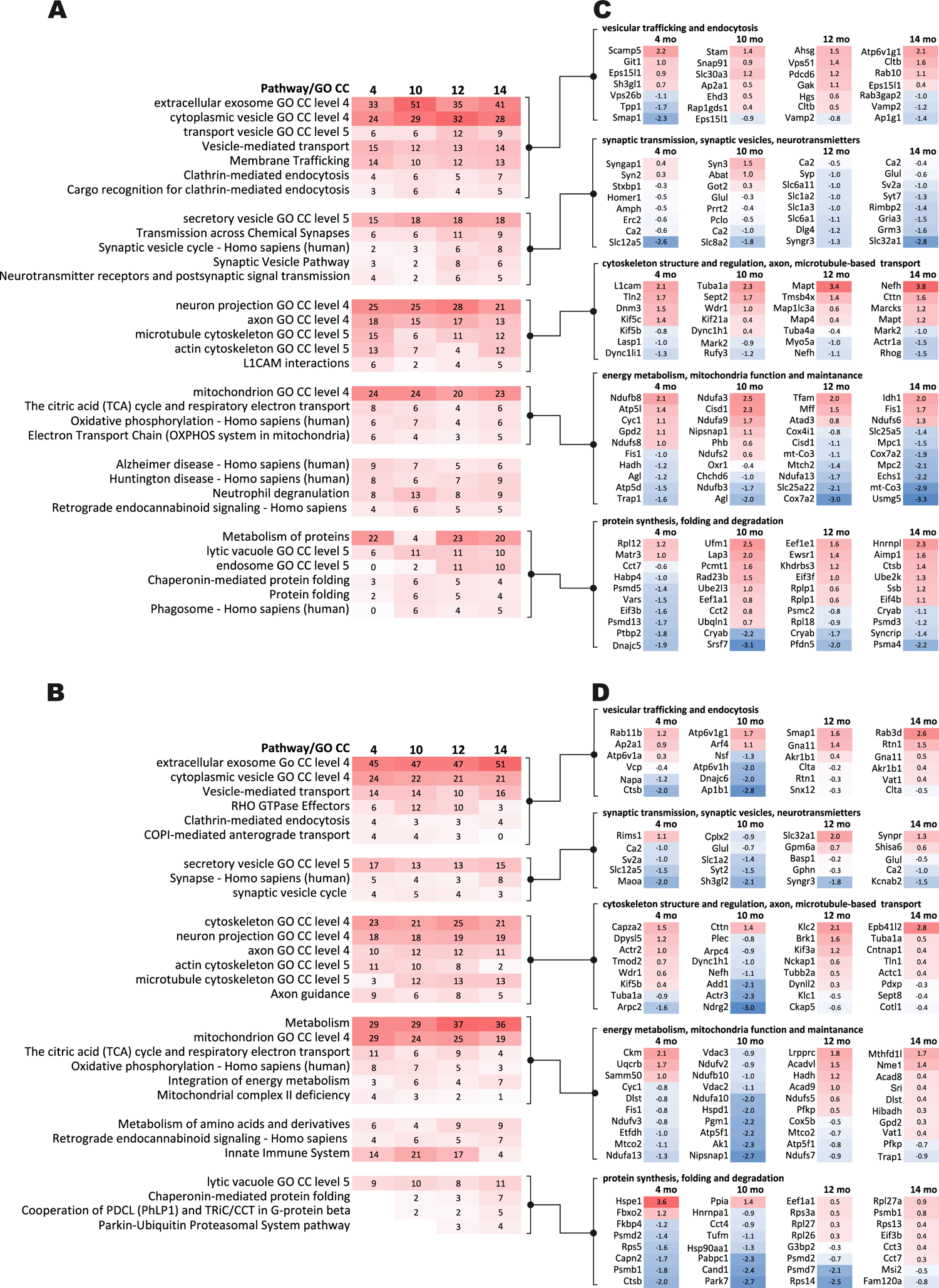
GO term and pathway analysis of dysregulated proteins identified in proteomics at behavioral milestones in the Ki91 SCA3/MJD mouse brain implicate vesicular transport, cytoskeleton, protein metabolism, and mitochondrial function. Bioinformatic analysis of biological pathways and subcellular localization of dysregulated proteins (p <0.05; two-sample t-test, n = 4 per genotype) during disease progression in the cerebral cortex (A) and cerebellum (B) was assessed using Consensus Path DB (CPDB; pathways p-value cutoff < 0.01; GO terms cellular component, p-value cutoff < 0.01, level 3, 4). The heatmaps present changes of pathways and subcellular localization involving dysregulated proteins (p <0.05; two-sample t-test) over the course of the disease in 4 tested ages (4, 10, 12, and 14-month-old) by showing what percentage of dysregulated proteins in each set is assigned to the particular pathway or subcellular region (numbers). The gradient of red color denotes a higher percentage of dysregulated proteins (more intense color) to a lower percentage of dysregulated proteins (less intense color). The 5 groups of pathways and GO terms arranged in blocks in panel (A) and (B) are linked to the corresponding lists of dysregulated proteins (p < 0.05; two-sample t-test) with the highest log2-FC in each tested age, assigned to 5 arbitrary categories in the cerebral cortex (C) and cerebellum (D). The categories were formed based on the analysis of pathways and GO terms and are as follows: (1) vesicular trafficking and endocytosis; (2) synaptic transmission, synaptic vesicles, neurotransmitters; (3) cytoskeleton structure and regulation, axon, microtubule-based transport; (4) energy metabolism, mitochondria function and maintenance and (5) protein synthesis, folding and degradation. The lists of dysregulated proteins are also presented as heatmaps with numbers representing log2-FC (C,D). The gradient of red color is for upregulated proteins, blue color for downregulated proteins.

The first category was related to vesicular transport and endocytosis. Cytoplasmic vesicles and extracellular exosomes were the localization for 24-32 % and 33-51% of dysregulated proteins in every dataset from the cortex (Fig. 4A) and similarly 21-24 % and 45-51% in the cerebellum (Fig. 4B). Moreover, vesicle-mediated transport and clathrin-mediated endocytosis are predicted to be disturbed in both the cerebral cortex (Vamp2, Cltb, Rab10; Fig. 4C) and cerebellum (Rab11b, Napa, Nsf, Rab3d; Fig. 4D).

The second category included dysregulated proteins involved in synaptic transmission. There was a relative increase in the number of altered proteins in older 12-14 month-old Ki91 mice in the cerebral cortex compared to younger ages (4-10 mo). For instance, pathways such as “transmission across chemical synapses” contained ∼ 6 % of dysregulated proteins in 4-10 mo vs ∼10 % in 12-14% and “synaptic vesicle cycle” ∼ 3% vs. 7%, respectively (Fig. 4A). Proteins related to SNARE complex and synaptic vesicles such as Syn3, Syp, Sv2a, and Syt7 (Fig. 4C) were dysregulated. Moreover, many proteins belong to the solute carrier (SLC) group of transporters, interestingly all downregulated (Fig. 4C). In general, most of the proteins with a function in the synapse showed decreased levels in the cerebral cortex of older 12-14-month-old Ki91 mice (Fig. 4C). In the cerebellum, dysregulated proteins of synaptic vesicles or synapses were less frequent compared to the cortex (“synaptic vesicle cycle” 3-5 % of dysregulated proteins; Fig. 4B) and included Sv2a, Syt2, and Kcnab2 (Fig. 4D). In both tested tissues, downregulation of Car2, Qdpr, and Glul, were detected in most of the tested ages (Fig. 4C, D), which we also validated using vacuum dot-blot assay (p<0.05; two-sample t-test; Suppl. Fig. 2). Glul, Car2, and Qdpr play a role in the metabolism of amino acids and neurotransmitters, and Qdpr level is slightly and gradually dropping down across ages (Suppl. Table 1, 2).

The third category was referring to altered proteins playing a role in regulating cytoskeletal structure and function. Dysregulated proteins localized to neuronal projections (axons and dendrites) constituted between 21 and 28% in the cortex and ∼18 % in the cerebellum (Fig. 4 A, B). Proteins assigned to cytoskeletal localization included actin (cortex: Actr1a; cerebellum: Arpc2, Actr3), microtubules (cortex: Tuba1a, Mapt) and neurofilaments (cortex and cerebellum: Nefh) (Fig. 4C, D). Disturbances of the cytoskeleton structure may lead to cellular transport defects, and there were many proteins directly regulating anterograde and retrograde axonal transport. For instance, in the cerebral cortex, dysregulated proteins involved in anterograde microtubule-based transport were Kif5c, Kif5b, and Kif21a and retrograde transport: Dnm3, Dync1li1, Dync1h1, and microtubule-associated proteins: Map1lc3a and Mark2 (Fig. 4C). In the cerebellum, proteins involved in the anterograde microtubule-based transport included Kif5b, Klc1, Klc2, and Kif3a, and in retrograde transport Dync1h1 and Dynll2 (Fig. 4D).

The fourth category involved proteins with altered levels involved in energy metabolism and mitochondria maintenance. Proteins predicted to localize in mitochondrion were between 20 to 24% and 19 to 29% in cortex and cerebellum, respectively (Fig. 4A, B). The top pathway involving dysregulated proteins in the Ki91 cerebellum was metabolism (Fig. 4B). Moreover, the number of proteins dysregulated in the cerebellum and associated with metabolism increases with age (4-10 mo: ∼ 29%; 12-14 mo: ∼ 36%; Fig. 4B). Proteins related to TCA and oxidative phosphorylation are dysregulated in the cerebral cortex (mitochondrial proteins Ndufb8 with; Ndufa3, Ndufs6, mt-Co3; Fig. 4C) and cerebellum (Ndufa13, Ndufa10, Ndufs7; Fig. 4D). There were also dysregulated proteins responsible for mitochondria fission (Fis1, Mff) and mitochondrial chaperons (Hspd1, Trap1) (Fig. 4C, D). Also, several proteins with altered levels mostly related to metabolism were linked to Alzheimer’s and Huntington disease in both the cerebellum and cerebral cortex (Fig. 4A, B).

The fifth category included dysregulated proteins involved in protein metabolism. In the case of the cerebral cortex, there are 4-23% dysregulated proteins involved in the metabolism of proteins (Fig. 4A). Noteworthy, the number of dysregulated proteins predicted to localize in lysosomes (6 % in 4 mo vs. ∼10 % in 10-14 mo), endosomes (2 % in 10 mo vs. ∼10 % in 12-14 mo), and phagosomes (4-6% in 10-14 mo) is higher in the cortex of older mouse (Fig. 4A, B). Similarly, the number of altered proteins involved in protein folding (2-3% in 10-12 mo vs. 5-7 % in 14 mo) and “Parkin-Ubiquitin Proteasomal System pathway” (3-4% in 12-14 mo) is increasing with age in the Ki91 cerebellum (Fig. 4B). Dysregulated proteins in this category involve various ribosomal subunits and translation factors (cortex: Rpl12, Eif3b; cerebellum: Rpl27a), proteasome subunits (cortex: Psmd13, Psma4; cerebellum: Psmb1, Psmd7) and chaperons (cortex: Dnajc5, Cryab, cerebellum: Hspe1, Park7) (Fig. 4C, D). Downregulation of Cryab, which was one of the most frequently identified proteins implicated in the pathology of neurodegenerative disease (Zhu and Reiser, 2018), was confirmed with vacuum dot blot assay (p < 0.05; two-sample t-test) in the cerebral cortex (10, 14 and 18 mo; Suppl. Fig. 2A) and cerebellum (18 mo; Suppl. Fig. 2B). Of note, most of the proteasome subunits and chaperons demonstrated decreased levels in both tested tissues (Fig. 4C, D).

The cellular origin of dysregulated proteins was investigated by searching for potential cellular markers of affected cell types in the SCA3 Ki91 brain using Dropviz (Saunders et al., 2018). In general, dysregulated proteins in sets from all ages originated from all main cellular types in the brain (neurons, oligodendrocytes, astrocytes). However, in the cerebellar cortex, we identified a relatively large group of dysregulated proteins originating from oligodendrocytes and polydendrocytes (Suppl. Fig. 3). The proteins included Cryab, Bcas1, Cnp, and Slc12a2, Car2, Odpr, Rhog, Gstp1, and Mbp (Mbp confirmed by dot blot assay (p <0.05; Suppl. Fig. 2A).

### 4. Proteomics correlative with SCA3 behavioral phenotype in 18-month-old Ki91 mice reveals dysregulated proteins involved in transport, cytoskeleton, synapse, and mitochondria

Our correlative proteomic approach aimed to discover dysregulated proteins in the brain of animals that demonstrated consistent behavioral phenotype to establish a more exact relationship between the protein biomarkers and the degree of SCA3 disease severity. Therefore, after the endpoint of behavioral studies (18 months), the animals were grouped by the intensity of phenotype, and 3 phenotype subgroups were established in the Ki91 cohort: severe, moderate, and mild (n = 4). The selection was based on score assessment on the 0-5 scale (Kruskal-Wallis test, p <0.05) (Fig. 5A). The scores 0 or 1 represented the “mild phenotype” group, score 2 or 3 was the “moderate phenotype” group, and 4 or 5 was the “severe phenotype” group. The cerebellum and cerebral cortex were collected from each group, and the tissues were subjected to proteomics (Suppl. Table 3; Suppl. Fig. 4 A, B). The analysis of dysregulated proteins revealed that protein levels in the moderate group were mostly in between the level of the severe and mild group, demonstrating a gradual pattern of protein dysregulation in Ki91 mice (Fig. 5B, C). Therefore in further data processing, we focused on the protein levels in severe and mild phenotype.

**Figure 5.**
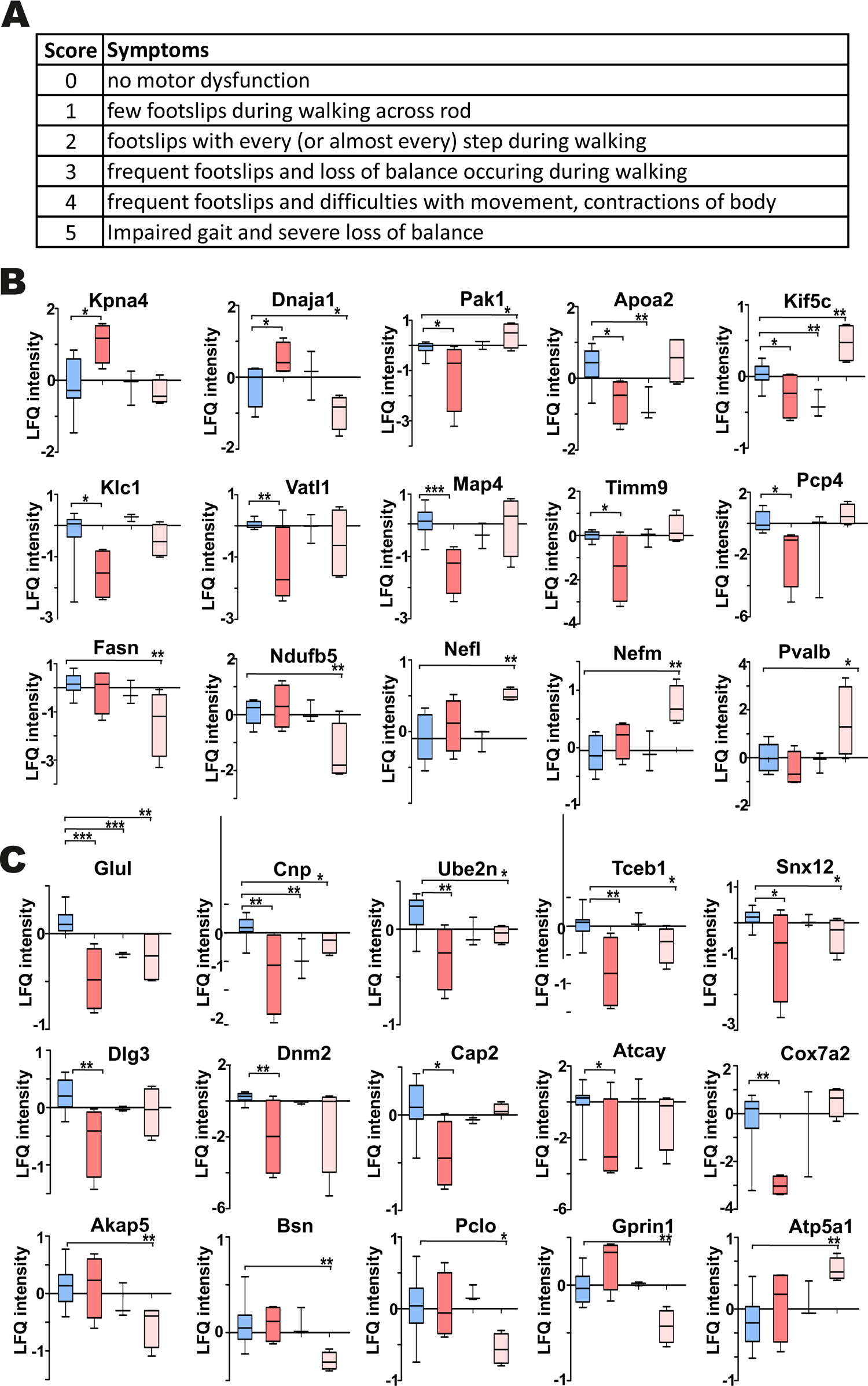
Correlative proteomics on brains of Ki91 SCA3/MJD mice demonstrate dysregulated proteins involved in intracellular transport in severe SCA3 phenotype and mitochondria, cytoskeleton, and synaptic vesicles in mild phenotype. To increase the correlation of behavior and proteomics, we selected 3 phenotype subgroups in the Ki91 cohort during behavioral testing of 18-month-old mice: severe, moderate, and mild (n =4). The selection was based on score assessment evaluating motor impairments in the 0-5 scale (Kruskal-Wallis test, p <0.05) (A). In the cerebellum derived from animals with a severe phenotype, there were upregulated (p < 0.05; two-sample t-test) heat shock protein Dnaja1 and a nuclear importin Kpna4 and downregulated proteins related to microtubule-based transport (Klc1, Kif5c, Vatl1, and Map4) and Purkinje cell protein (Pcp4) (B). In the cerebellum of animals with mild phenotype, neurofilament proteins were upregulated (Nefl, Nefm) together with Pvalb, Pak1, Kif5c (p < 0.05; two-sample t-test) (B). Mitochondrial proteins were downregulated in both severe (Timm9) and mild phenotypes (Ndufb5 and Fasn) (B). In the cerebral cortex of animals with severe phenotype proteins related to intracellular transport and cytoskeleton (Snx12, Dlg3, Dnm2, Cap2, and Atcay) and mitochondrial protein Cox7a2 were downregulated (p < 0.05; two-sample t-test) (C). Animals with mild phenotype displayed lower levels of proteins associated with synaptic vesicles and post-synaptic density (Akap5, Bsn, Pclo, Gprin1) (C). The blue color is for WT, and the gradient of pink color is for Ki91, the most intense pink color is a severe phenotype, and the palest pink color is for the mild phenotype. two-sample t-test (*P < 0.05, **P < 0.01, ***P < 0.001), n = 36, n = 4 per phenotype, error bars: SEM.

We found that in the cerebellum of the Ki91 animals with the severe phenotype (n = 4), there were heat shock protein Dnaja1 and a nuclear importin Kpna4 upregulated compared to WT animals (WT; n = 10) (Fig. 5B). Another member of the importin family, Kpna3, was previously shown to control the nuclear localization of ataxin-3 (Sowa et al., 2018). There were also 5 upregulated proteins in the mild phenotype (n = 4) – Pvalb, Pak1, Kif5c, and neurofilaments (Nefl and Nefm). Pak1, a kinase implicated in vesicle-mediated transport and regulation of cytoskeleton dynamics, and Kif5c involved in microtubule-based transport, were also downregulated in severe phenotype (Fig. 5B). Moreover, in the severe group, there were also other downregulated proteins playing a pivotal role in transport along the microtubule (Klc1, Kif5c, Vatl1, and Map4) and notably Purkinje cell protein 4 (Pcp4), which was not dysregulated in mild phenotype (Fig. 5B). Noteworthy, mitochondrial proteins were downregulated in both severe and mild phenotypes. Mitochondrial chaperone Timm9 showed lower levels in the “severe” group, while Ndufb5 and Fasn were downregulated in the “mild” group (Fig. 5B).

In the cerebellar cortex, the only upregulated protein was a subunit of mitochondrial ATP synthase, Atp5a1 (Fig. 5C). Proteins commonly downregulated in both severe and mild phenotype (Fig. 5C) included Glul, already identified in the proteomics performed parallel to behavioral milestones and myelin protein Cnp, ubiquitinating enzyme Ube2n, transcription factor Tceb1, and Snx12 involved in intracellular trafficking. In the mild group only, there were downregulated proteins associated with synaptic vesicles and post-synaptic density (Akap5, Bsn, Pclo, Gprin1; Fig. 5C)). Whereas in a “severe” group, there were downregulated proteins related to intracellular transport and cytoskeleton (Snx12, Dlg3, Dnm2, Cap2, and Atcay), and mitochondrial protein Cox7a2 (Fig. 5C).

Altogether, we identified a set of proteins involved in intracellular and microtubule-based transport downregulated in the cerebellum and cerebral cortex of Ki91 mice displaying severe phenotype. Downregulated proteins in the “mild” phenotype were related to mitochondria in the cerebellum and synapse in the cerebral cortex. Proteins upregulated in the cerebellum were associated with cytoskeleton and neurofilaments in the “mild” group. Cellular processes and compartments associated with dysregulated proteins in “severe” phenotype were related to mitochondria, intracellular and microtubule-based transport, synaptic vesicles, and the cytoskeleton. Most of these processes and compartments were also identified in our approach parallel to behavioral milestones. An essential neuronal structure, which connected those terms, is the axon, which requires high energy production, highly efficient trafficking of cargoes toward its end, and is a location of synaptic vesicle release. Therefore, we further decided to go to the next level of neuronal complexity in our proteomic investigation and selectively explore proteins dysregulated in the SCA3 axons.

### 5. Targeted axonal proteomics indicates the abnormal localization of proteins between axons and soma such as cytoskeletal, vesicular proteins and highly increased translation machinery in Ki91 SCA3 neurons

Our next aim was to define with high precision the specific processes which highly contribute to axonal dysfunction in SCA3 already at a very early level of neuronal maturation stage, where no neuronal damage takes place, and therefore no secondary post-symptomatic events occur. Therefore, we performed the analysis of enriched or decreased proteins in axons of the cerebellar and cortical neurons (Fig. 6, 7). Neurons from wildtype and Ki91 E18 embryos (cortical neurons) and P5 pups (cerebellar neurons) were cultured on a 1-μm porous filter in Boyden chambers (n = 4 per genotype, each replicate represented neuron culture isolated from a different set of embryos or pups) (Fig. 6A). The axons were growing through the pores to the other side of the filter, which enabled the separate collection of the neuronal somatodendritic and axonal compartments (Suppl. Fig. 5 A, B). Both somatodendritic and axonal protein extracts were subjected to MS/MS proteomics, and the “raw levels” for each protein for both compartments were determined (Suppl. Table 4). To normalize the protein levels within fractions originating from neurons on the same filter, we calculated the protein level ratio between the axonal and somatodendritic compartments for each WT and Ki91 sample/filter (Suppl. Table 4). Such normalization has proved very solid since the comparison of soma/axon ratios within the single genotype (WT or Ki91) demonstrated that the most significant difference was the abundance of nuclear proteins, such as histones, and nuclear enzymes characteristic for the somatodendritic compartment (ratio axon to soma < −3; Suppl. Table 4). Coherently, the PCA graph demonstrates a clear separation between axonal and somatodendritic samples (Suppl. Fig. 5 C, D).

**Figure 6.**
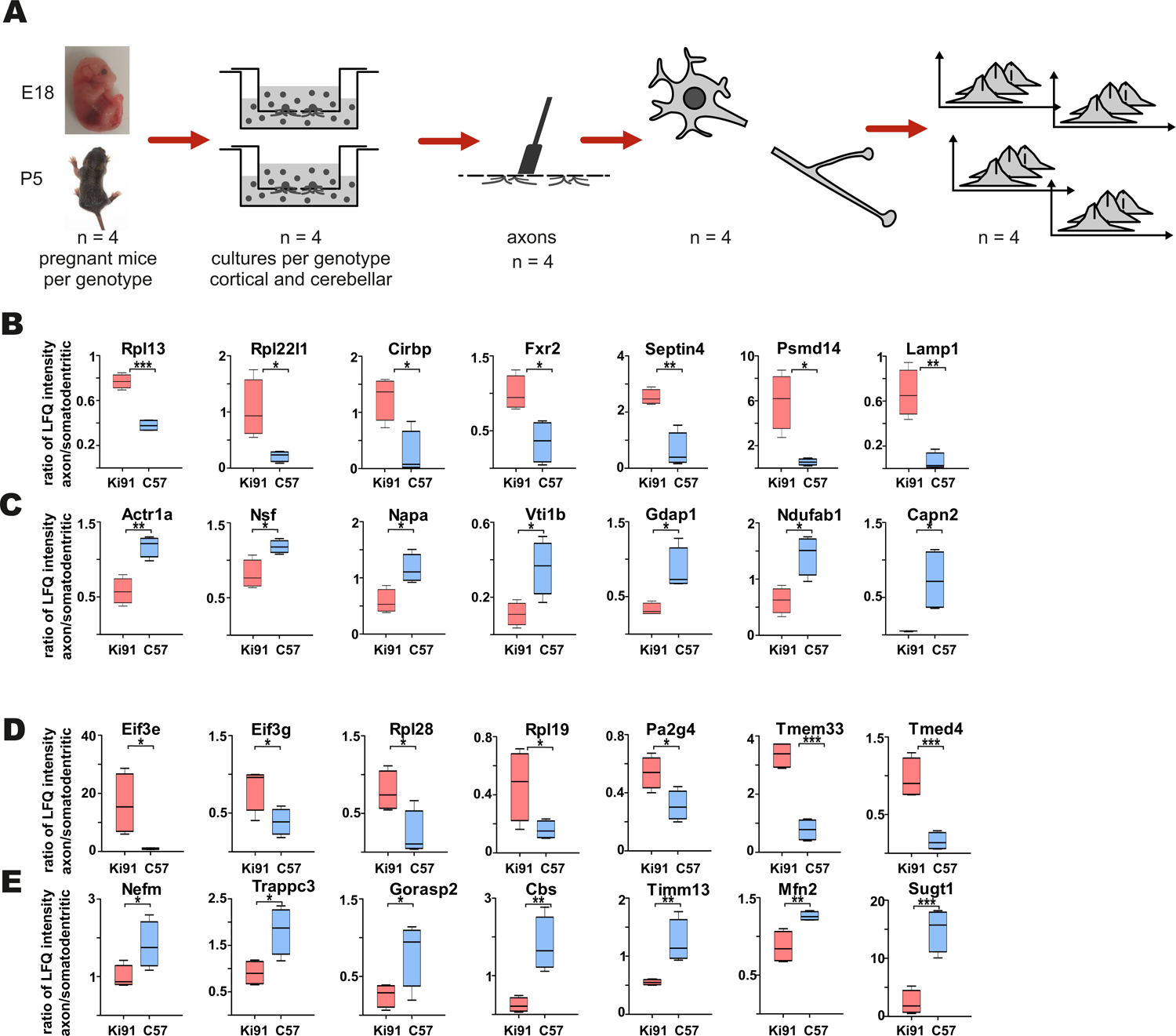
Targeted axonal proteomics reveals sets of proteins enriched or depleted in the Ki91 SCA3/MJD cerebellar and cortical axons. Identification of proteins differentially localized in SCA3 neuronal compartments was performed by isolating two neuronal fractions (axonal and somatodendritic) from the same culture (Boyden chambers) obtained from E18 embryo (cortical) or P5 pups (cerebellar) – derived neurons. In brief, neurons were seeded on the inner side of the Boyden chamber inserts with a 1-μm porous filter. After the period of neuronal differentiation and axonal growth through and towards the outer side of the insert membrane, both the inner side of the insert (somatodendritic) and the outer side (axonal) were scraped, and the fractions were collected for MS/MS analysis (A). Graphs present proteins with abnormal localization between axon and soma with the highest ratio between those compartments (p < 0.05; two-sample t-test) in the primary cerebellar (B, C) and cortical Ki91 neurons (D, E). In the P5-derived Ki91 cerebellar axons, there were enriched proteins related to ribosomes (Rpl13, Rpl22l1), RNA-binding (Cirbp, Fxr2), and proteasome or lysosome (Psmd14, Lamp1) (B). Proteins depleted in the Ki91 cerebellar axons involved important modulators of vesicular transport (Actr1a, Nsf, Napa, Vti1b) (C). In the E18-derived Ki91 cortical axons, there were enriched translation factors (Eif3e, Eif3g), ribosome subunits (Rpl28, Rpl19), RNA-binding (Pa2g4), and ER and Golgi (Tmed4, Tmem33) (D). Proteins that were depleted in the Ki91 cortical axons included neurofilament (Nefm), related to intracellular trafficking (Trappc3, Gorasp2), and mitochondria (Timm13, Mfn2) (E). two-sample t-test (*P < 0.05, **P < 0.01, ***P < 0.001), n = 36, n = 4 per phenotype, error bars: SEM.

Subsequently, we compared protein level ratios between WT and Ki91 SCA3/MJD neurons to calculate the fold change and estimate if the protein was enriched or depleted in axons. Then, we searched for proteins that displayed statistically significant differential fold change between WT and Ki91 group (p < 0.05; two-sample t-test; Suppl. Table 4).

In cerebellar neurons, there were 34 proteins enriched (ratio axon/soma > 1, p < 0.05; two-sample t-test; n=4) and 41 proteins depleted (ratio axon/soma < 1, p < 0.05; two-sample t-test; n=4) in Ki91 vs WT axons (Fig 6 B; Suppl. Table 5, 6). There were 27 and 20 such proteins in cortical neurons, respectively (Fig. 6 C; Suppl. Table 7, 8). The analysis of proteins in both cerebellar and cortical Ki91 axons revealed a common functional characteristic. The majority of the axon-enriched proteins in SCA3 samples were related to the upregulation of protein production and degradation (Fig. 6B, C; 7A, B; Suppl. Tables 6, 8). Strongly upregulated was the translation machinery, in particular the initiation step and nonsense-mediated decay together with the RNA binding proteins, including Cirbp (FC = 4.92; Fig. 6B), which stabilizes mRNAs involved in cell survival during stress (Liao et al., 2017). Highly upregulated were the proteasome subunits and chaperones enriched in cerebellar Ki91 axons (Psmd14, Psma5, and Hspa8; Suppl. Table 6) and cortical Ki91 axons (Sel1l, Cct2; Suppl. Table 8), which altogether might be related to the neuronal response to cellular stress and misfolded proteins in SCA.

In contrast, proteins that were reduced in Ki91 axons comparing to WT were in the majority related to the actin cytoskeleton and vesicle-mediated transport, which may indicate the axonal structural dysfunction (Fig 6 B,C; 7A, B; Suppl. Table 5, 7). In Ki91 cerebellar axons, there were reduced levels of 13 proteins related to cytoskeleton, including 8 proteins regulating actin function (Fig. 6B, 7A; Suppl. Table 5). The actin cytoskeleton is involved not only in maintaining the proper structure of the axon but also plays a pivotal role in the short-range transport, selection of cargoes targeted to the axon tip, formation of synapses, and scaffold for the mitochondria in the growth cone (Kevenaar and Hoogenraad, 2015). Along this line, we identified 8 mitochondrial proteins with a lower ratio axon to soma in cerebellar SCA3 samples as compared to WT (Fig. 6B, 7A; Suppl. Table 5). Altogether, this may indicate the disrupted delivery of mitochondria to the proper axon area due to the deregulated function of the actin cytoskeleton. Moreover, there are also proteins with a reduced level in cerebellar Ki91 axons directly related to intracellular trafficking (Actr1a, Ank2, Arf6, Nsf, Rhob), synaptic/SNARE vesicles (Sv2b, Napa, Vti1b), and COPI-mediated transport from ER to Golgi (Napa, Actr1a, Nsf, Sptan1, Sptbn, Ank2) (Fig. 6B, 7A; Suppl. Table 5).

In cortical Ki91 axons, there were generally fewer proteins identified with lower ratio axons to the soma (Fig. 7B, Suppl. Table 7). However, there were also proteins related to mitochondria, transport, and cytoskeleton, including neurofilament medium-chain (Nefm), playing an essential role in axonal function (Fig. 6C, Fig. 7B, Suppl. Table 7).

**Figure 7.**
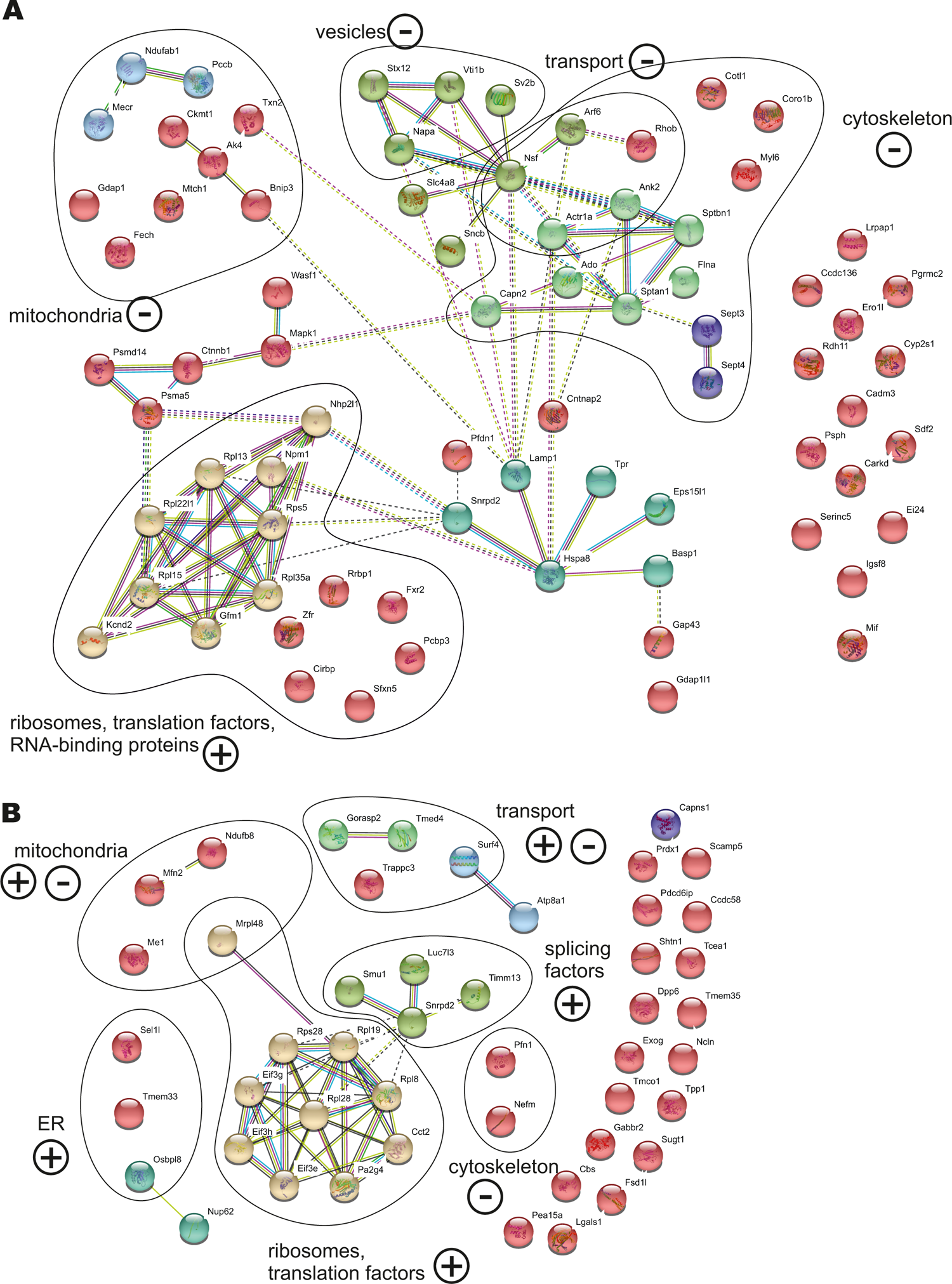
Abnormal localization between axon and soma in the cerebellar and cortical neurons of Ki91 SCA3/MJD build an intensely interconnected network of proteins of translation machinery, vesicles and transport machinery, cytoskeletal and mitochondrial proteins. The network of proteins displaying altered ratio between axon and soma in cerebellar (A) and cortical neurons (B) of Ki91 mice were generated using String database and clustering (https://string-db.org/). Number of proteins enriched in cerebellar axons n = 34 (ratio axon/soma > 1, p < 0.05; two-sample t-test); reduced in cerebellar axons n = 41 (ratio axon/soma < 1, p < 0.05; two-sample t-test); enriched in cortical axons n = 27 (ratio axon/soma > 1, p < 0.05; two-sample t-test); reduced in cortical axons n = 20 (ratio axon/soma < 1, p < 0.05; two-sample t-test), 4 biological replicates per genotype. Distinct colors denote clusters of proteins generated by the String algorithm, which grouped functionally associated proteins into specific sets, such as translational machinery (ribosomes, translation factors), cytoskeletal-modulators, and vesicular proteins. Functional groups are marked and named based on GO annotations (BP, M Level 3, 4, and 5; q <0.01) using CPDB or GO term search if not available in CPDB. “+” and “–“ denote for enrichment or depletion of protein groups in axons based on F.C. of the majority of proteins in the group. Blue and pink edges states for known interactions, and green, red, and dark blue for predicted interactions, and black and purple for others.

### 6. Impairment of mitochondrial metabolic potential in Ki91 SCA3/MJD primary cerebellar neurons

SCA3 patients show decreased BMI, and similarly, SCA3 Ki91 mice fail to gain weight. Moreover, our proteomic approaches identified mitochondria and energy metabolism failure in Ki91. The direct investigation of energy metabolism in presymptomatic young SCA3 neurons was not performed previously; therefore, it is unknown if the mitochondrial phenotype occurs early in SCA3 and if mitochondrial dysfunction is a primary contributor to the SCA3 pathology or only the secondary result of putative mice and patient dysphagia. To address whether energy metabolism is disturbed in Ki91 neurons, we performed Cell Energy Phenotype Test with Seahorse XFp analyzer. The assay was performed on primary cerebellar neurons in DIV3, 11, 18, and 21 (n=6 per genotype and stage; Fig. 8 A - C). The oxygen consumption rate (OCR) and extracellular acidification rate (ECAR) were measured over time at baseline and stressed conditions (injection of oligomycin and FCCP mixture) (Suppl. Fig. 6). Then, the metabolic potential of neurons was evaluated by calculating changes after stress conditions relative to baseline. The test demonstrated a significant increase in the ECAR metabolic potential in both Ki91 and WT neurons in every tested stage (p < 0.0001; two-sample t-test; Fig. 8A, B). Likewise, an increase of the OCR metabolic potential was observed in WT neurons and Ki91 neurons, but only in DIV11 and 21 (p < 0.05; two-sample t-test; Fig. 8A, B). Furthermore, under stressed conditions, the levels of ECAR were diminished in Ki91 neurons compared to WT in DIV3, 18, and 21 (p < 0.001; two-sample t-test; Fig. 8A, C). Interestingly, both baseline and stressed OCR was increased in Ki91 neurons compared to WT in DIV 3 and 11 (p < 0.0001; two-sample t-test; DIV11 baseline p = 0.13; Fig. 8A, C), while in later stages, the trend was reversed and in DIV18 and 21 it was significantly decreased (p < 0.01; two-sample t-test; Fig. 8A, C).

**Figure 8.**
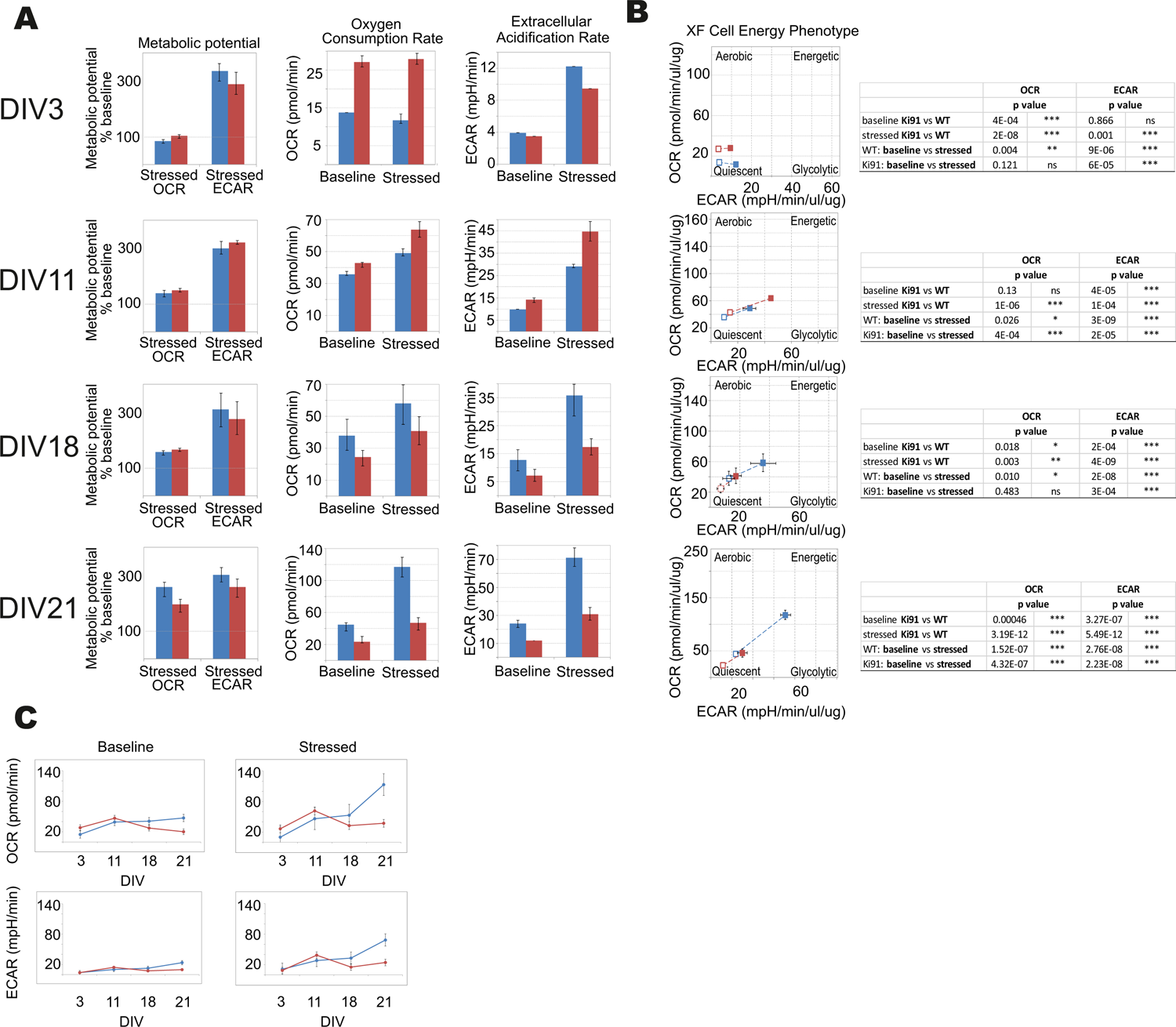
The energy metabolism measured by Seahorse XFp Cell Energy Phenotype Test is impaired in the P5 cerebellar neurons of Ki91 SCA3/MJD. The assays were performed with cerebellar neurons at DIV3, 11, 18, and 21. The rate of mitochondrial respiration (OCR, oxygen consumption rate) and glycolysis (ECAR, extracellular acidification rate) were measured under baseline and stressed conditions, which were evoked by specific stressors: 1 μM of oligomycin and 1 μM of FCCP. The bioenergetic parameters are shown for OCR (pmol/min), ECAR (mpH/min), and metabolic potential ((stressed OCR or ECAR/baseline OCR or ECAR) × 100%)) in the panel (A). In DIV3 and DIV11, OCR in neurons is elevated in Ki91 compared to WT (p<0.001; two-sample t-test), but in later stages (DIV21), both OCR and ECAR are severely decreased in Ki91 neurons (p<0.001; two-sample t-test) under both baseline and stressed conditions (A). WT cerebellar neurons transform from the quiescent phenotype in DIV3 towards energetic in DIV21, but Ki91 does not undergo such transformation (B). The profiles of OCR and ECAR during tested developmental stages *in vitro* are presented in panel (C). The blue color is for WT neurons; red is for Ki91 neurons. Experiments were performed in n = 6 per genotype and stage. Two-sample t-test (*P < 0.05, **P < 0.01, ***P < 0.001), error bars: SEM.

Overall, these data demonstrate that the potential for energy production is impaired in cerebellar Ki91 neurons. The glycolysis rate shows fluctuations in both baseline and stressed conditions. On the other hand, the mitochondrial respiration shows a coherent pattern, which starts with an increase under both tested conditions, following by a decrease. The study demonstrates that energy deficit occurs early in SCA3 neurons, and therefore it is not a secondary event resulting from previous neuronal damage.

### 7. Assessment of vesicle state in neurites of Ki91 SCA3/MJD cerebellar neurons

All proteomic approaches collectively demonstrated altered levels of proteins building cellular vesicles or regulating intracellular vesicular trafficking. Therefore, our next goal was to assess a state of vesicles using the functional assay based on the labeling of neuronal vesicles using Rab7, a vesicular protein commonly occurring in early and late endosomes, lysosomes, and other types of vesicles and in vesicles that are undergoing active transport in axons and dendrites. Therefore we transfected mRFP-tagged Rab7 to SCA3 primary cerebellar neurons (neuron cultures isolated from 3 different sets of P5 pups, 5 independent transfections, with 5-10 neurons analyzed each time; Fig. 9 A).

**Figure 9.**
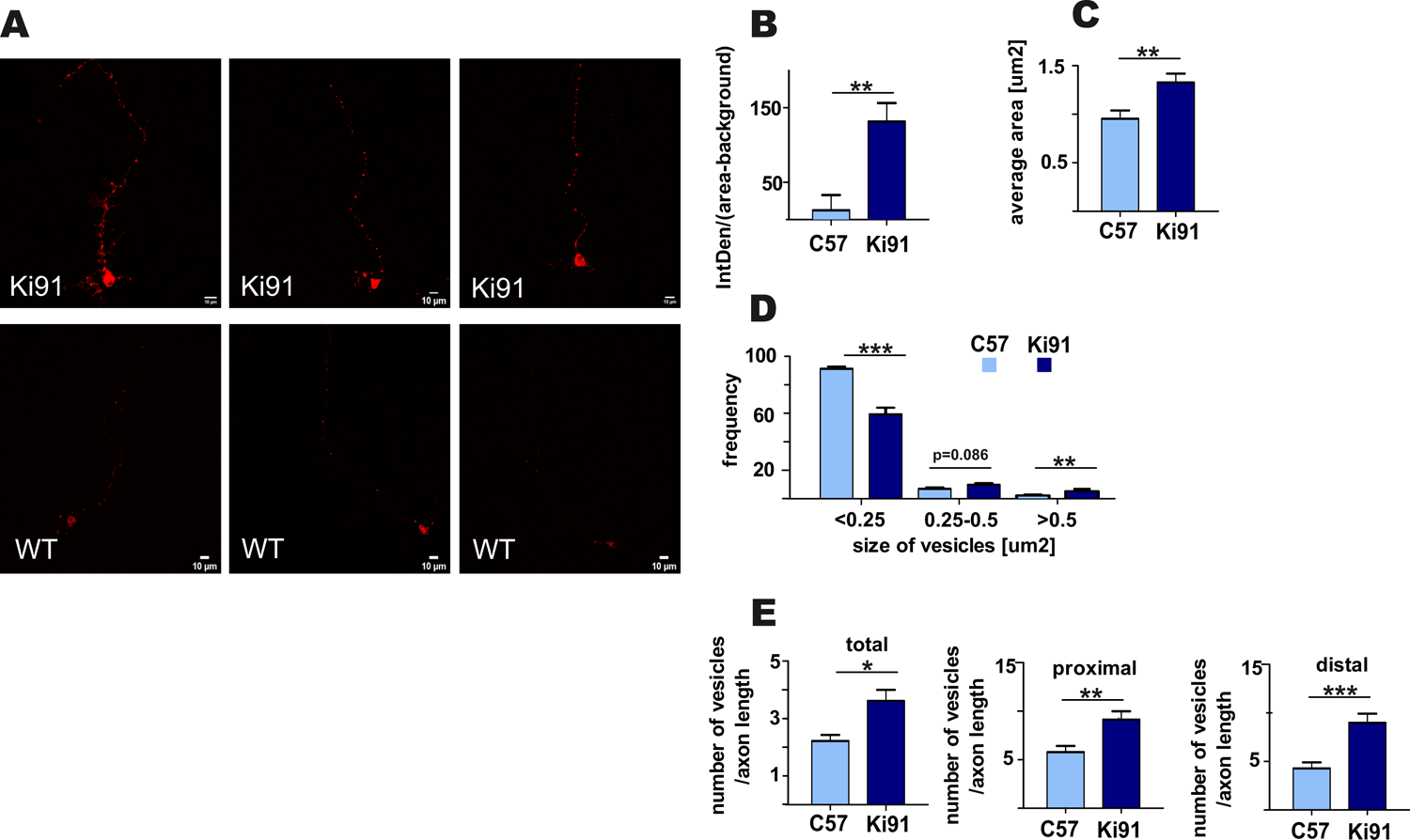
Accumulation and enlargement of vesicles in Ki91 SCA3/MJD P5 cerebellar neurons. To investigate if the consensus vesicular and transport process identified by various types of proteomics in Ki91 impacts vesicle appearance and number, we transfected the primary cerebellar neurons (DIV8-9) with RFP-labeled Rab7. The Rab7 was selected as a useful vesicle tag since it is often recruited to several types of vesicles, including late endosomes and lysosomes (red). Representative images show fluorescently tagged vesicles in cerebellar neurons from Ki91 mice (upper panel) and WT neurons (lower panel) (A). The sizes and the number of tagged vesicles were quantified using ImageJ. The mean intensity of fluorescence was much higher in Ki91 cerebellar neurons (p<0.01; two-sample t-test) (B). The average area of fluorescently tagged vesicles was 1.33 μm2 (n = 45) for Ki91, 0.95 μm2 (n =34) for WT (p<0.01; two-sample t-test) (C). The size distribution of tagged vesicles in Ki91 cerebellar neurons showed a shift from smaller to larger sizes in comparison to WT (p<0.001; two-sample t-test) (D). The number of vesicles per axon length was higher in Ki91 neurons, also measured separately for both proximal and distal parts (defined as 1/8 of the axon length starting either from the axon hillock or axon tip) (E). The measurements were from 5 independent experiments, with 5-10 cells analyzed each time. Scale bars: 10 μm. Two-sample t-test (*P < 0.05, **P < 0.01, ***P < 0.001), error bars: SEM.

We found that the labeling of vesicles with fluorescently tagged Rab7 was much more intensive in Ki91 than in WT neurons (Fig. 9 B). Closer investigation revealed that the average area of vesicles was significantly (t-test, p < 0.01) increased by 40% in Ki91 as compared to WT neurons (Fig. 9 C). Distribution of vesicle size showed that Ki91 neurons had a lower number of small vesicles (<0.25 μm2) and a higher number of large vesicles (>0.5 μm2) (t-test, p < 0.01; Fig. 9 D). We did not detect any differences in the shape of vesicles. However, there was a significant increase in the number of vesicles normalized to the length of the measured axon (t-test, p < 0.05; Fig. 9 E). To assess whether Rab7-loaded vesicles accumulate in either the proximal or distal region of the axon, we also calculated the number of vesicles in those regions, arbitrarily defined as 1/8 of the axon length starting either from the axon hillock or axon tip. There was a significant increase in the number of vesicles in the proximal region (t-test, p < 0.01) by 58% in Ki91 as compared to WT neurons and in the distal part (t-test, p < 0.001) by 110% (Fig. 9 E).

### 8. Ki91 SCA3/MJD neurons show increased phosphorylation of axon-specific neurofilaments, indicating the stress of the axonal cytoskeleton

The proteomic approaches and, in particular, the axonal proteomics demonstrated that several classes of dysregulated proteins in SCA3 are located in the axonal compartment. We also found the ataxin-3-positive inclusions along axons in the cerebellum by co-staining with axon-specific marker smi-312 antibody (axon-specific p-Nef). The phosphorylation state of axonal neurofilaments reflects the health of the neuronal cytoskeleton. Therefore we further explored whether the structure and phosphorylation status of neurofilaments are disturbed in the SCA3 model. We assessed the phosphorylated and non-phosphorylated neurofilaments (axon-specific p-Nef and neuronal Nef; Fig 10) on immunostained sagittal brain slices from 12-month-old Ki91 and WT mice using. Increased phosphorylation of neurofilaments is often a sign of defective transport and maintenance also related to mitochondria. We found that both p-Nef levels (SMI-312 antibody) and non-phosphorylated Nef levels (SMI-32 antibody) were significantly increased in the cerebellum of SCA3 mice compared to WT animals (t-test, p < 0.001; n=3 and 5 pictures per genotype; Fig. 10 A-D). In detail, we found increased levels of Nef in soma, dendrites, and axons of Purkinje cells (t-test, p < 0.01; Fig. 10 A, C). The increased level of axon-specific p-Nef was detected in axons of the molecular layer and axons of the granular layer and white matter (t-test, p < 0.001; 10 B, D). The altered phosphorylation of axons in the SCA3 Ki91 model may be related to disturbed protein and mitochondria localization and their transport in axons.

**Figure 10.**
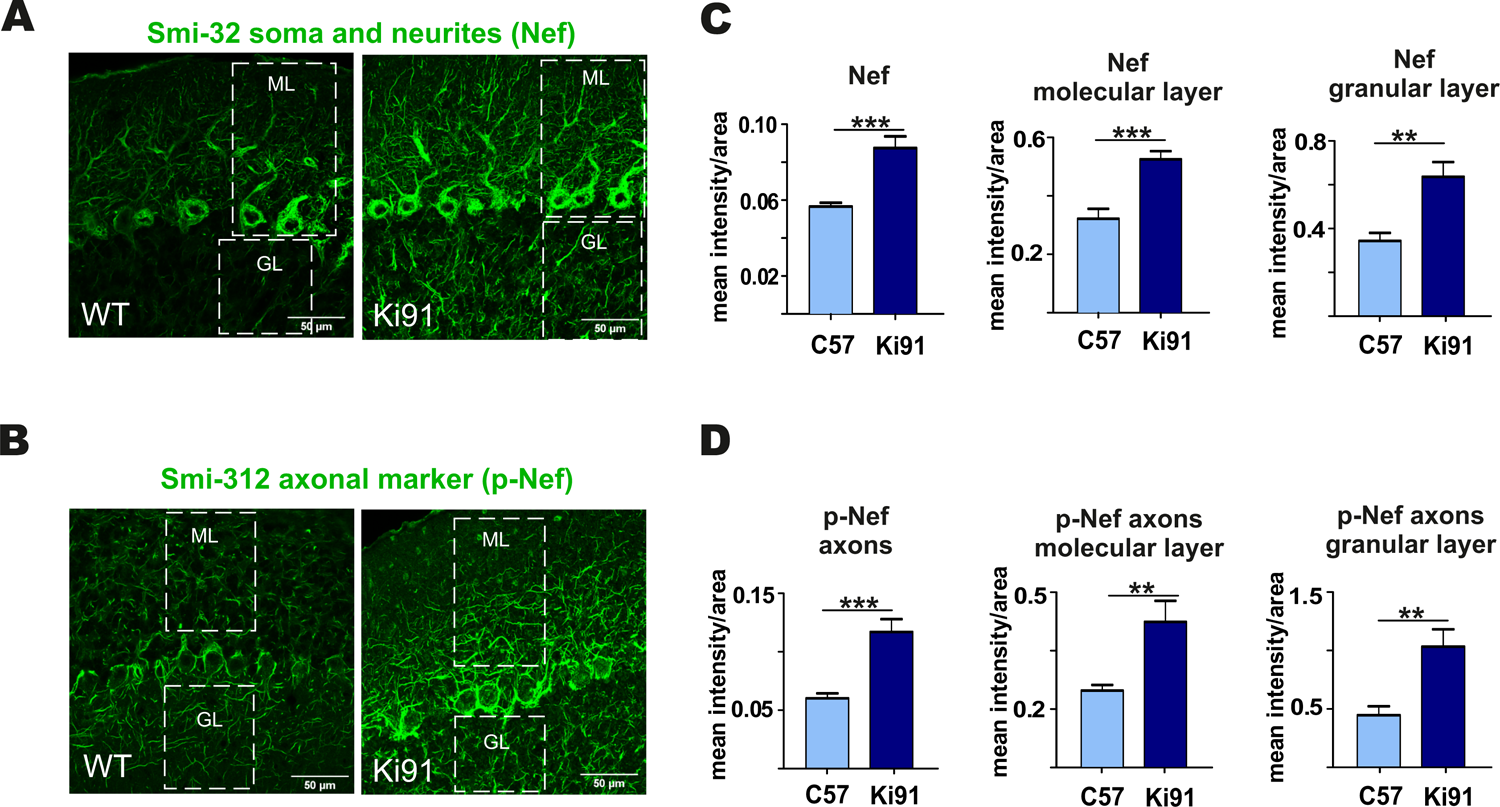
Accumulation of neurofilaments in the cerebellar neurons of Ki91 SCA3/MJD brain. Immunofluorescent staining of the cerebellar sections from 12-month-old mice evaluating the condition of neuronal cytoskeleton and phosphorylation state of neurofilaments in axons. The staining show accumulation of phosphorylated neurofilaments in SCA3 cerebellar axons, which is often a sign of axonal stress. In addition, the non-phosforylated pan-neuronal neurofilaments also show increased staining intensity indicating altered cytoskeleton in SCA3 neurons. Scale bars: 50 μm (A–B). Non-phosphorylated heavy neurofilaments (Nef) detected with SMI-32 antibody (pan-neuronal) are presented in panel (A). Phosphorylated neurofilaments H and M detected with SMI-312 (p-Nef; axon-specific) are shown in panel (B). White boxes inside pictures (A) and (B) mark the area, which was used for measuring the mean intensity of the fluorescence comprising the molecular and granular layer of the cerebellum. Measurement of mean intensity of fluorescence using ImageJ and statistics (p<0.05; two-sample t-test) performed for sections as a whole, and for both layers separately is presented on the graphs (C, D). Axon-specific SMI-312-positive neurofilaments were highly phosphorylated in both layers in the cerebellum of the Ki91 animals (molecular layer, p<0.001; granular layer, p<0.01; two-sample t-test) (B, D). Moreover, the level of dephosphorylated neurofilaments was also elevated in both cerebellar layers of the Ki91 (p<0.01; two-sample t-test) (A, C). N = 3 biological replicates and 10 confocal images per genotype were collected.

## Discussion

The protease/deubiquitinase activity of ataxin-3 suggests its widespread influence on cellular proteome and changes of various brain protein levels resulting from mutant ataxin-3 and SCA3. Surprisingly, so far, very little is known about the proteome in the SCA3 brain, and it may be one of the reasons for the incomplete understanding of the disease mechanism and delay in finding a cure. Here, we aimed to investigate the brain profile of altered proteins in the SCA3 knock-in model to unveil a consensus set of most critical molecular programs governing the disease and identify protein biomarkers. In detail, we aimed to investigate the proteome using three different approaches: at the *in vivo* level, at the cellular compartment level, and in relation to disease progression and severity. Once the consensus set of altered candidate processes was established, we aimed to test if we can identify their physiological fingerprints by simple, functional assays, preferably using primary presymptomatic neurons.

We first performed a longitudinal behavioral study between 4 months and 18 months of age of Ki91 (Switonski et al., 2015) to establish SCA3 behavioral and phenotypic milestones suitable for the investigation of brain proteome. The first milestone in symptomatic Ki91 mice was the reduced body weight gain identified in 4-month-old animals. Statistically significant gait ataxia at 12-month-old Ki91 was the second milestone, and the last milestone was the severe motor disturbances and progression of other symptoms in 18-month-old animals. Altogether, homozygous Ki91 represents a model of SCA3 progression and gradation of symptoms observed in SCA3 patients (Coutinho and Andrade, 1978; Jacobi et al., 2015; Pulido-Valdeolivas et al., 2016; Coarelli et al., 2018).

In the next step, we determined a range of affected brain areas by the MRI and by detection of ataxin-3-positive inclusions using advanced, symptomatic Ki91 animals at 18 months of age. The inclusions were present in almost all brain areas of both the posterior and frontal brain parts. The intra-nuclear inclusions dominated in the cerebellum, mainly in the white matter, DCN, and in pons and midbrain; however, they were also present in the cerebral cortex, hippocampus, and striatum. Importantly, we have seen highly abundant ataxin-3-positive inclusions in SMI-312-positive cerebellar axons in white matter. The intra-axonal inclusions were also previously detected in nerves and various tracts in SCA3/MJD patients (Seidel et al., 2010). Also, the changes observed by brain MRI in Ki91 were not localized to one structure, but the atrophy was spreading on several brain regions, including frontal parts, cerebellum, and brainstem. Moreover, brain areas rich in axons such as the corpus callosum are atrophied in Ki91. We could not consistently measure the atrophy of the cerebellum by MRI (see material and methods); however, the cerebellum in Ki91 is affected, showing the most significant number of inclusions and cerebellar features of Ki91 were demonstrated by us previously (Switonski et al., 2015).

Therefore, we have selected two representative brain regions, the cerebellum representing the more posterior part of the brain and the cerebral cortex representing frontal brain areas, for further proteomic studies. Our first proteomic approach corresponded to Ki91 behavioral milestones where we selected tissue from animals at the age of 4 months (lack of weight gain), 10 months (mice on the verge of behavioral phenotype), 12 months (motor decline statistically significant), and 14 months (an additional symptomatic age). The second proteomic approach, called “correlative,” was aimed to identify dysregulated proteins and processes in mouse subjects with carefully characterized and matching phenotype severity. The first two proteomic approaches offered insights into the disease pathogenesis at the level of tissue and disease stage; however, the perspective of cellular level and cellular compartments were still missing in both approaches. Therefore we employed a third, targeted proteomics approach where we separately assessed proteins in somatodendritic and axonal compartments of primary neurons isolated from embryos or pups. An additional advantage of such primary neuronal SCA3 culture is the possibility of validation if processes discovered in two other approaches may occur early in the disease development.

The first pathogenic process, which was consensus among the proteomic approaches in our work, is the disturbed energy metabolism. We demonstrated many dysregulated proteins related to the mitochondrial compartment, which corresponds with the reduced body weight gain in Ki91. Notably, the BMI of SCA3 patients is significantly lower, and BMI decrease correlated with faster disease progression (Diallo et al., 2017; Yang et al., 2018). The energy metabolism was the most common pathway, and the proteins related to metabolism were dysregulated already in 4-month-old presymptomatic Ki91 mice. For instance, many OXPHOS-related proteins, such as Nduf subunits, Aco2, and mt-Co proteins and several metabolic proteins such as Gpd2, Cs, Gpi, Pkm, Wdr1, Sdha, Ddx1, and Aldh1l1 demonstrated altered levels and were previously identified in AD, HD and SCA1 (Sorolla et al., 2008; Sánchez et al., 2016; Lunnon et al., 2017). Similarly, the correlative approach also revealed downregulated mitochondrial proteins such as mitochondrial chaperone Timm9, Ndufb5, and Fasn in both “mild” and ‘severe” phenotype, which suggest that disturbances of energy production is the primary pathology of SCA3. Several mitochondrial proteins were also depleted in axons compared to the somatodendritic compartment in SCA3 neurons, which may indicate the disrupted delivery of mitochondria to nerve terminals. The most frequently dysregulated protein, which we propose as one of a SCA3 biomarker candidate in both tested brain regions, is carbonic anhydrase 2 (Ca2). Ca2 is linked to a mechanism controlling an increased transport of lactate, which is an important energy source for neurons (Stridh et al., 2012; Karus et al., 2015).

Since many proteomic changes pointed out pathways related to energy metabolism, our first functional assay measured mitochondrial respiration and glycolysis rate in cultured cerebellar Ki91 neurons kept for DIV 3, 11, 18, and 21, which reflected the culture differentiating from young to more mature neurons. Commonly, the typical culture of such neurons and also our WT neurons represented a developing dependency on OXPHOS since mature neurons rely on oxidative phosphorylation to meet energy demands. In turn, SCA3 neurons demonstrated a lack of increase in OXPHOS dependency and even a slight drop in oxygen consumption during neuronal maturation. SCA3 neurons under stress reached their maximal oxygen consumption rate already as young as DIV 11 as compared to WT neurons, which reached the maximum at DIV 21. At DIV 3 and 11, SCA3 neuronal progenitors show higher overall oxygen consumption; therefore, at this early stage, the neurons may already be in deficiency of energy, or its production is aberrant. In the case of older SCA3 neurons with highly developed processes, a dramatic difference in energy metabolism compared to WT neurons can also be related to the failure of energy production in axons.

Recent discoveries emphasize the role of degeneration of “wiring” between brain structures and axonal deficits as one of the main pathogenic events in neurodegenerative disorders (Sánchez-Valle et al., 2016; Ferrari Bardile et al., 2019; Casella et al., 2020). Also, in SCA3, the axonal defects and white matter abnormalities are prominent disease symptoms (de Rezende et al., 2015; Farrar et al., 2016; Lu et al., 2017). Our proteomic approaches jointly demonstrated that the largest population of proteins with altered levels was involved in vesicular transport and endocytosis as a second consensus molecular process candidate in SCA3. Vesicular trafficking is crucial for axon maintenance, synaptic signaling, releasing, and recycling of neurotransmitters. Our proteomic data demonstrate highly downregulated levels of proteins related to vesicle transport or trans-synaptic signaling (Nrgn, Sh3gl1, Vamp2, Syngap1, Bsn, Syt7, Homer1, Rims1, Pclo, Erc2; SNARE complex). Several vesicular Rab proteins were also dysregulated, such as Rab11b, Rab3d, and Rab10. Other Rab proteins were also dysregulated in SCA3/MJD patients (Sittler et al., 2018). Besides, there were dysregulated proteins related to protein metabolism localized to endosomes, exosomes, and lysosomes. Many of these proteins are involved in neurodegenerative diseases such as AD and HD (Skotte et al., 2018; Mendonça et al., 2019), and their loss brings significant neurological consequences (Yoo et al., 2001; Santos et al., 2017; Salpietro et al., 2019; Falck et al., 2020). Moreover, Qdpr (human homolog: DHPR) and Glul proteins responsible for the metabolism of neurotransmitters are progressively downregulated throughout the SCA3 course in the Ki91 cerebellum and cortex. Both proteins are essential for proper CNS function (Breuer et al., 2019; Kurosaki et al., 2019; Zhou et al., 2019).

The 3rd consensus SCA3 candidate process was termed by us “the regulation of cytoskeletal structure and function” with dysregulated proteins localizing to neuronal processes. Coherently with vesicle trafficking, our proteomic results also revealed dysregulation of neurofilament proteins (NFs), such as heavy chain NF-H in both the cerebral cortex and cerebellum. Disturbed energy metabolism in neurons is often related to defects in the cellular cytoskeleton and phosphorylation state of NF-H, in particular, resulting in aberrant axonal transport of mitochondria (Jung et al., 2000) (Shea and Chan, 2008). Furthermore, proteomic data contained dysregulated proteins pivotal for microtubular cytoskeleton stability and axon organization. Dysregulated proteins were also involved in anterograde and retrograde microtubule-based transport (Kif5c, Kif5b, and Kif21a, Klc1, Klc2, Kif3a; Dnm3, Dync1li1, Dync1h1, Dynll2), in particular in “severe” phenotype, which may suggest that disruption of axonal transport contributes to increased severity of SCA3 symptoms. Among the microtubular proteins, Tau (Gao et al., 2018) was elevated in the cerebral cortex of symptomatic Ki91 mice, RhoG (Zinn et al., 2019), and dysregulated group of proteins that are also enriched in oligodendrocytes such as Cryab protein stabilizing cytoskeleton and previously proposed as a component of therapeutic strategy (Ghosh et al., 2007; Houck and Clark, 2010) (Cox and Ecroyd, 2017; Zhu and Reiser, 2018). Dysregulation of Cryab, RhoG, Mbp, and Marcks indicates that oligodendrocytic processes and myelination contribute to axon dysfunction in SCA3 (Ishikawa et al., 2002; Guimarães et al., 2013; Zhang et al., 2017; Rezende et al., 2018). Interestingly, several oligodendrocytic marker proteins dysregulated in our proteomic studies, such as Plp, Qdpr, Mbp, and Mog were predicted to localize in extracellular vesicles (EVs). EVs are often released by oligodendrocytes and deliver trophic support for axons but may also transfer pathogenic triggers (Krämer-Albers et al., 2007) (Porro et al., 2019).

The investigation of two candidate consensus processes related to vesicle transport and neuronal cytoskeleton required a targeted proteomic approach focused on axons. The proteins that were deficient in SCA3 axons were associated with vesicles, axonal transport, and mitochondria. Depleted proteins suggesting altered transport along the axon in Ki91 included proteins interacting or forming dynactin complex, required for mitochondria and endosome/lysosome and synaptic vesicles transport along the axons, such as Actr1a and Ank2 (Holleran et al., 1996; Moughamian et al., 2013; Lorenzo et al., 2014). Besides, there were also essential cytoskeletal proteins involved in regulating axonal transport and actin dynamics, Arf6 and its interactor RhoB, targeted by Arf6 to endosomes (Eva et al., 2017; Zaoui et al., 2019). Components of SNAREs responsible for vesicular trafficking were also depleted in SCA3 axons, such as Napa (a.k.a. α-SNAP), Vti1b, and Nsf. Lower levels of Napa, Nsf, and Actr1a were identified by the “parallel” proteomic approach in the cerebellum of Ki91 mice. Of note, Nsf is the ATPase indispensable for membrane fusion and for regenerating active SNAREs. Nsf depletion results in the arrest of membrane trafficking and, consequently, neuronal damage and death (Stenbeck, 1998; Emperador-Melero et al., 2018; Yuan et al., 2018). In particular, disturbed axonal transport may affect delivery and proper localization of mitochondria since we detected several essential mitochondrial proteins with diminished levels in Ki91 axons.

In addition to defective energy metabolism, all of our proteomic approaches identified the vesicular processes and cytoskeletal structure and regulation processes in the SCA3 Ki91 brain. In addition to vesicular and cytoskeletal localization, most dysregulated proteins identified in proteomics were predicted to localize in axons. Therefore we designed two simple assays to detect functional fingerprints of vesicular and cytoskeletal phenotypes in neurons. We decided to load the neuronal vesicles with fluorescent Rab7 protein as the universal, external marker often occurring in a variety of vesicle species. We were able to detect the accumulation of vesicles in cell bodies and in the processes of SCA3 neurons. Moreover, the vesicles were enlarged and shown intensive fluorescence in SCA3 neuronal processes. The second assay was based on immunostaining detection of axonal cytoskeleton phosphorylation to assess a putative pathogenic process related to the axonal cytoskeleton. We used the SMI-312 antibody, which is axon-specific marker (phosphorylated NFs), and detected highly increased phosphorylation in the 12-month-old Ki91 cerebellum compared to WT cerebellum. Non-phosphorylated NFs (SMI-32; pan-neuronal staining) were also increased in the SCA3 KI91 cerebellum. Accumulation and aggregation of phosphorylated neurofilaments in axons is a biomarker indicating axonal damage, which may disturb the transport of vesicles and mitochondria. Moreover, in neurodegenerative disorders and SCA3, the NFs could serve as a biomarker (Li et al., 2019).

Lastly, axonal proteomics in Ki91 has also demonstrated highly enriched protein translation machinery represented by ribosomal proteins, various RNA-binding proteins, and translation initiation factors (Suppl. Table 5, 7). Protein synthetic machinery is intensively transported to the axons in response to cellular stress (Baleriola and Hengst, 2015). We detected the signs of cellular stress by identifying elevated levels of Tpp1 and Tmem33 proteins, which are playing a role in unfolded protein response (UPR) (Sakabe et al., 2015; Mukherjee et al., 2019). Along this line, protein chaperons and proteasome subunits (Cct2, Pmsd14) were highly enriched in SCA3 Ki91 axons. Moreover, ribosomal and RNA-binding proteins are also compounds of RNP granules (mRNPs) or stress granules, both holding mRNAs translationally repressed (Sossin and DesGroseillers, 2006; El Fatimy et al., 2016). One of the crucial components of the granules is Fxr2 (Chyung et al., 2018), which is highly enriched in Ki91 cerebellar axons. Interestingly, vesicular platforms in axons were recently identified as carriers of translation machinery, mRNPs, and structure responsible for axonal mitochondria maintenance (Cioni et al., 2019). Furthermore, it was also demonstrated that protein inclusions specifically impair the transport of endosomes and autophagosomes (Volpicelli-Daley et al., 2014; Guo et al., 2020). In addition, locally active translation machinery and local energy production are long known phenomena in axons since it provides critical substrates even to the most distal part of the neuronal terminals. However, in the case of accumulation of the polyQ proteins, such an organization of protein synthesis in long processes may accelerate neuronal dysfunction. In such a case, upregulated translation machinery may be a “vicious cycle” for the accumulation of polyQ proteins in axons, gradually aggravating the disease and leading to profound neuronal dysfunction.

## Conclusions

SCA3 molecular disease mechanism is insufficiently explored. Ataxin-3 is ubiquitin protease, but very little is known about the broader influence of mutant protein on other proteins in the SCA3 brain. The missing knowledge about the proteome may be one of the reasons for the incomplete understanding of the disease mechanism and lack of cure. Therefore, we present the first global model picture of SCA3 disease progression on the protein level in the brain, in embryonic SCA3 axons, combined with an *in vivo* behavioral study and neuropathology. We propose the most relevant pathogenic processes in SCA3 involving disturbed energy metabolism, impairment of vesicular system, structure of cell and axon, and altered protein homeostasis. We demonstrate that the consensus processes identified in the SCA3 Ki91 brains show their functional signs in the primary pre-symptomatic neurons, and therefore may be the first steps of pathogenesis. Our data have demonstrated for the first time that one of the essential pathogenic aspects of SCA3 is the faulty localization of proteins between axons and soma. Further signs of affected SCA3 axons are vesicle accumulation and increased neurofilament phosphorylation. The important novel finding is the highly increased axonal localization of proteins involved in the translation machinery in SCA3. It is possible that local mutant protein production in axons may accelerate neuronal dysfunction; therefore, upregulated translation machinery in axons may aggravate the disease. Moreover, we identified a variety of molecular signatures related to mitochondria in the SCA3/MJD Ki91 model, and we show that the earliest global process which occurs already in the neonatal neurons is the OXPHOS and glycolysis deficits. For the first time, we demonstrate that the young developing SCA3 neurons fail to undergo the transition from glycolytic to OXPHOS phenotype typical for normal neurons. Summarizing, we revealed that the SCA3 disease mechanism is related to the broad influence of mutant ataxin-3 on the brain, neuronal and axonal proteome, and disruption of the proper localization of axonal and somatodendritic proteins and organelles such as vesicles, translation machinery, and mitochondria (Fig. 11).

**Figure 11.**
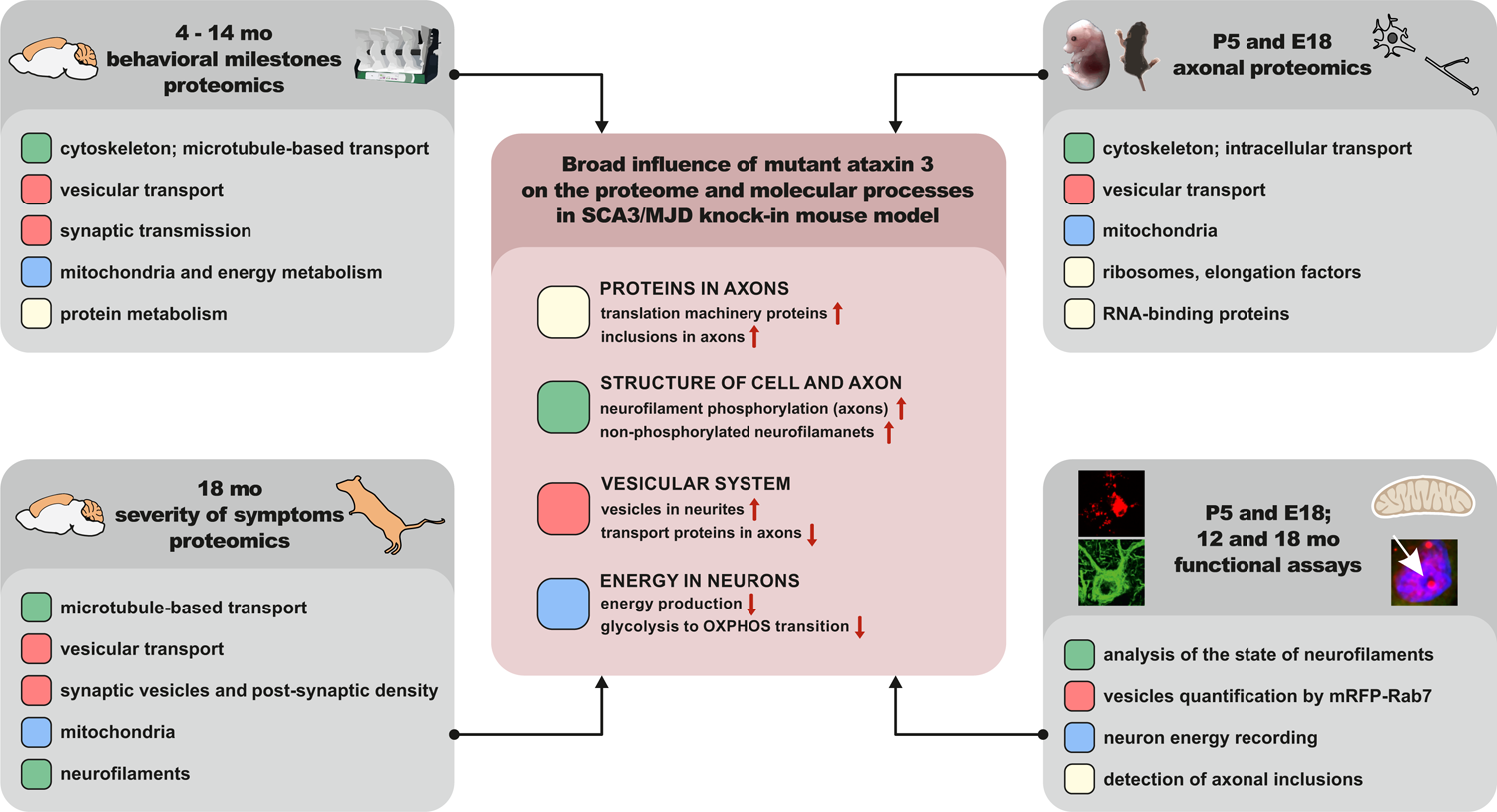
The graphical summary of broad influence of mutant ataxin 3 on the proteome and molecular processes in SCA3/MJD knock-in mouse model

## Material and Methods

### Animals

Maintaining and breeding were performed at standard conditions with an 18/6-h light/dark cycle and water and food ad libitum. No differences in food or water intake between Ki91 and WT mice were observed. The animals were marked using numerical ear tags (National Band & Tag Company, Newport, USA). The stress level of the animals was minimized throughout all procedures and animal handling. The animal experimentation and handling were approved and monitored by the Local Ethical Commission for Animal Experiments in Poznan. The animals were sacrificed according to AVMA Guidelines for the Euthanasia of Animals by placing them in the programmable CO2 chamber (Rothacher Medical, Heitenried, Switzerland). Homozygous SCA3 Ki91 animals contained between 103 and 132 CAG repeats on a single mutant ataxin3 allele. SCA3 Ki91 mice (C57BL/6 background) and age-matched wild-type littermates bred in-house were used for behavioral tests (n=36) and collecting brain tissues for proteomic analysis (n=32, and n=21, after behavioral tests), validation of proteomics (n=32), MRI (n=12) and immunostaining (n=8, after behavioral tests and n=6 from a separate cohort of 12-month-old animals). The cerebellum and cerebral cortex for proteomic analysis were collected from 4, 10, 12, 14, and 18 month-old animals, which corresponded to the timepoints of behavioral testing. An experimental group consisted of 4 mutant mice vs. 4 wildtype littermates (proteomics: 4, 10, 12, and 14 moth-old and immunostaining), and 11 mutant mice vs. 10 wildtype littermates (proteomics: 18 moth-old).

### Behavioral studies

Mice were trained and tested for motor deficits, starting from 4-month-old animals, and consequently every 2 months, until the age of 18 moth-old in a cohort of 18 mutant mice vs. 18 wildtype littermates. Tests included accelerating rotarod (4 – 40 rpm in 9.5 min), elevated beam walk (diameter of rods: 35 mm, 28 mm, 21 mm, 17 mm, 9 mm) as previously described (Switonski et al., 2015), and additionally in 18-month-old animals open field test in which locomotor activity and zone preferences in the experimental cage (50 cm2) were analyzed and measured during 10 minutes. Each test consisted of 1 training day (T) and 3 consecutive days of measurement. We also performed scoring tests designed for evaluation of the ataxia phenotype in mouse models (Guyenet et al., 2010), wire test for evaluating muscle strength, and footprint for gait ataxia. For gait analysis, the forelimbs of the mice were stained with black and hindlimbs with red non-toxic paint. Mice were placed on a sheet of paper inside a tunnel (80 × 10 cm). Only the middle part of a series of steps was measured for the distance between two steps (stride length), for- and hindlimb base, and overlapping between for- and hindlimb. All mice were weighed during each testing session. Graphing and statistics were performed on PrismVR software (San Diego, CA, USA), using ANOVA with Bonferroni post-hoc test.

### Magnetic Resonance Imaging and Image analysis

After PFA perfusion of the mice, brains (n=6 per genotype) were removed and stained by a one-week soaking in Gadolinium solution (Dotarem©, Guerbet, France) in PBS at 2.5 mM. This protocol enhances the signal- and contrast-to-noise ratios on MR images of fixed brains (Dhenain et al. 2006). Ex vivo MRI experiments were performed on a horizontal 11.7 T Bruker scanner (Bruker, Ettlingen, Germany). A quadrature cryoprobe (Bruker, Ettlingen, Germany) was used for radiofrequency transmission and reception. Diffusion Tensor Imaging (DTI) data were acquired using Echo Planar Imaging sequence (Resolution= 100×100×200 µm3, TR= 3000 ms, TE= 28 ms, 60 directions). To preserve their integrity and avoid deformations, brains were kept inside skulls for ex vivo MRI experiments. Anatomical images were co-registered in the same space to create a study template. Transformations to match the study template to an atlas composed of a high-resolution template and a map of region’s labels adapted from the Allen mouse brain Atlas (Lein et al. 2007) were calculated. Finally, transformations to match the study template and labels to each anatomic image were calculated. The automated segmentation pipeline was performed using an in-house python library (Sammba-MRI, https://github.com/sammba-mri/sammba-mri).

### Protein sample preparation

Protein extraction was performed as previously described (Wiatr et al., 2019). Mouse brain tissues were homogenized with a Mixer Mill MM400 (Retch, Haan, Germany), followed by a threefold cycle of freezing and thawing and bath sonication (3×3-minutes). Protein concentration was estimated using Quant kit (GE Healthcare, Chicago, Il, USA). Ten-microgram aliquots of proteins were diluted with 15 µl of 50 mM NH4HCO3 and reduced with 5.6 mM DTT for 5 min at 95°C. Samples were then alkylated with 5 mM iodoacetamide for 20 min in the dark at RT. Subsequently, the proteins were digested with 0.2 µg of sequencing-grade trypsin (Promega, Madison, WI, USA) overnight at 37°C.

### Liquid chromatography coupled to tandem mass spectrometry (LC-MS/MS)

The analysis of the proteome was performed with the use of Dionex UltiMate 3000 RSLC nanoLC system connected to the Q Exactive Orbitrap mass spectrometer (Thermo Fisher Scientific, Waltham, MA, USA). Peptides derived from in-solution digestion were separated on a reverse-phase Acclaim PepMap RSLC nanoViper C18 column (75µm × 25cm, 2µm granulation) using acetonitrile gradient (from 4 to 60%, in 0.1% formic acid) at 30°C and a flow rate of 300nL/min (for 230 min). The spectrometer was operated in data-dependent MS/MS mode with survey scans acquired at a resolution of 70,000 at m/z 200 in MS mode and 17,500 at m/z 200 in MS2 mode. Spectra were recorded in the scanning range of 300– 2000 m/z in the positive ion mode. Higher energy collisional dissociation (HCD) ion fragmentation was performed with normalized collision energies set to 25. Swiss-Prot mouse database was used for protein identification with a precision tolerance set to 10ppm for peptide masses and 0.08 Da for fragment ion masses. Raw data obtained for each dataset were processed by MaxQuant 1.5.3.30 version for protein identification and quantification. Protein was considered as successfully identified if the Andromeda search engine found at least two peptides per protein, and a peptide score reached the significance threshold FDR=0.01.

Obtained data were exported to Perseus software ver. 1.6.1.3 (part of MaxQuant package; Munich, Germany). Numeric data were transformed to a logarithmic scale, and each sample was annotated with its group affiliation. Proteins only identified by site, reverse database hits, and contaminants were removed from the results. Next, data were filtered based on valid values for proteins. Proteins that contained valid values in 75% of samples in at least one group were included as valid hits. Before statistical analysis, normalization of data was performed by subtracting the median from each value in a row. A two-sample t-test was performed on analyzed sample data with p-value < 0.05 was considered as significant, and differentiating proteins were normalized using the Z-score algorithm for the hierarchical clustering of data.

### Bioinformatic analysis of proteomic data

Proteins were grouped using the Consensus Path Database (CPDB) according to the pathways (pathway enrichment p-value cutoff < 0.01), GO term by molecular function, biological process, and cellular component (GO term B, MF level 5, CC level 4 and 5, p-value cutoff <0.01). CPDB integrates data from 30 different resources, including such databases as Reactome, KEGG, and BioCarta (Kamburov et al., 2013). Next, common pathways and GO terms for 4 datasets separately in both tissues (4, 10, 12, and 14 months of age) containing the highest relative number of dysregulated proteins and p-values were selected. The relative number of dysregulated proteins was calculated as a number of dysregulated proteins involved in the pathway or GO term divided by the total number of dysregulated proteins in a given dataset (%). For each analysis, the names of genes corresponding to the names of dysregulated proteins were used. In the separate analysis, we performed identification of cell types in the brain, which are affected by pathogenesis with the use of the Dropviz tool (Saunders et al., 2018) for the cerebellum and cerebral cortex. Significantly dysregulated (p< 0.05) proteins identified from label-free proteomic and experiments were included together into two tissue groups based on origin from the cerebellum and cortex. Relative expression is presented in Dropviz as the number of transcripts per 100 000 in the cluster. The analysis of proteins displaying altered ratio between axon and soma in cerebellar and cortical neurons of Ki91 mice was performed using the String database (https://string-db.org/).and clustering tool (MCL clustering, inflation parameter = 6).

### Vacuum dot immunoblot

Before use in the dot blot assay, each antibody was validated with Western blot assay (WB). WB was performed as previously described (Wiatr et al., 2019). Cerebellum and cortex samples were harvested from 16 homozygous Ki91 animals and 16 wildtype littermates; n = 4 mice per tested age (4, 10, 14, 18 months of age) and per each genotype. The protein concentration was estimated using a Pierce BCA protein assay kit (Thermo Fisher Scientific, Waltham, MA, USA). The nitrocellulose membrane and Whatman filters (all GE Healthcare, Chicago, Il, USA) were washed 2X with Milli-Q H_2_0 and 2X with Tris-Buffered Saline (TBS) by vacuum filtration. 2.5 ug of each protein sample in 50 μL of PB lysis buffer with PMSF cocktail inhibitor (Sigma-Aldrich, St. Louis, MO, USA) was spotted on nitrocellulose membrane and washed 2X with TBS by vacuum filtration, and then allowed to air-dry. The blots were stained with Ponceau S solution and blocked with 5% nonfat milk in PBS/0.05% Tween 20 for 1 h at RT and subsequently incubated at 4 °C overnight with the following primary antibodies: rabbit anti-MBP (1:1000; Cell Signaling, Danvers, MA, USA), mouse anti-CRYAB (1:1000; Developmental Studies Hybridoma Bank, Iowa City, IA, USA), mouse anti-GLUL (1:1000; BioLegend, San Diego, CA, USA), rabbit anti-CA2 (1:2000; ProteinTech, Rosemont, IL, USA), rabbit anti-QDPR (1:2000; ProteinTech, Rosemont, IL, USA). The blots were probed with the respective HRP-conjugated secondary antibody (anti-rabbit or anti-mouse, 1:2000; Jackson Immuno Research, Suffolk, UK). The immunoreaction was detected using the ECL substrate (Thermo Fisher Scientific, Waltham, MA, USA).

### Primary neuron culture and transfection

Primary neuron cultures were derived from Ki91 and C57 mice according to AVMA Guidelines for the Euthanasia of Animals. Cortices were dissected from E18 mouse embryos and cerebella from P5 pups. Cortical neurons were dissociated by trypsin (Merck, Darmstadt, Germany) diluted 10×, and cerebellar neurons were dissociated by trypsin diluted 5x. Cells were washed 2 times in disassociation medium consisting of HBSS (Thermo Fisher Scientific, Waltham, MA, USA), supplemented with 0.1% glucose, 1x penicillin-streptomycin (Thermo Fisher Scientific, Waltham, MA, USA), then 2 times in plating medium consisting of Dulbecco’s modified Eagle medium (DMEM), 1x CTS GlutaMAX-I supplement, 1x penicillin-streptomycin (all Thermo Fisher Scientific, Waltham, MA). Clumps of tissue debris were allowed to settle to the bottom of a 15 ml tube, and the supernatant containing dissociated cells was centrifuged for 3 min at 1,300 rpm. Cells were seeded onto coverslips, tissue culture inserts (Greiner Bio-One GmbH, Kremsmünster, Austria), or Seahorse XFp Cell Culture Miniplates (Agilent Technologies, Santa Clara, CA, USA) coated with poly-D-lysine (Thermo Fisher Scientific, Waltham, MA, USA). Neurons were maintained in conditioned Neurobasal medium (Thermo Fisher Scientific, Waltham, MA, USA) supplemented with 2% B27 supplement, 1x CTS GlutaMAX-I supplement, 1x penicillin-streptomycin (all Thermo Fisher Scientific, Waltham, MA, USA), 1x apo-transferrin (Sigma-Aldrich, St. Louis, MO, USA), and N2 (0,005mg/ml insulin, 0,0161 mg/ml putrescin, 30nM Na-Selenite, 51nM T3, 20nM progesterone) in a humidified incubator with 5% O2 and 5% CO2 in air at 37 °C. The maintenance medium was replenished with half-feed changes every 2-3 days.

For transfection, cells were seeded at a density of 2 × 10^5^ cells/well onto coverslips in 24-well culture plates. Neurons were transfected at DIV 3-4 with mRFP-Rab7 (0.8 μg DNA per well) using Lipofectamine 2000 (Thermo Fisher Scientific, Waltham, MA, USA) followed by fixation (4% PFA, 15 min in RT) and confocal imaging at DIV 7-11 from 3 biological replicates and 5 independent transfections. An approximately 10%-15% transfection efficiency was achieved.

### Isolation of axons

Dissociated neurons (cortical and cerebellar) were seeded at a density of 1 × 10^6^ cells/well onto tissue culture inserts containing porous membrane (1 µm pores; Greiner Bio-One GmbH, Kremsmünster, Austria) in a 6-well cell culture plates. The bottom compartment medium was supplemented with 15 ng/ml of BDNF (PeproTech, London, UK) so that it could act as a chemoattractant for axons growing through the insert membrane. Axonal fraction and the fraction containing neuronal body with dendrites were isolated by carefully detaching the cellular contents from the inner and outer insert membrane surface with a cell scraper in TAEB buffer (Merck, Darmstadt, Germany) four times, alternating the direction by 90° each time. Axons were collected after 11 days in culture because, after DIV14, dendrites, which grow at a slower rate, are also detected on the other side of the porous membrane. Before isolation of the axon part, the inner membrane surface was scrubbed with a cotton-tipped applicator, and the membranes were cut out from the insert. To examine whether the separation of axons from the somatodendritic part was correctly performed, we also immunostained culture filters isolated from the Boyden chamber before and after scraping off the cell bodies, with immunofluorescence assay (β-III-tubulin, 1:500; Burlington, MA, USA) and nuclear dye Hoechst (Suppl. Fig. 5 A, B). The presence of axons on the outer site of the membrane was examined, and the purity of the obtained axonal fraction was evaluated with nuclear and axonal markers (Suppl. Table 3). The axon isolation procedures were adapted from (Unsain et al., 2014).

### Seahorse XFp Cell Energy Phenotype Test

Seahorse XFp Cell Energy Phenotype assays (Seahorse, Agilent Technologies, Santa Clara, CA, USA) were performed according to the manufacturer’s instructions on an XFp instrument. Dissociated cerebellar neurons were seeded at a density of 5300 or 12 500 cells/well onto culture mini plates with the protocol described in the section “Primary neuron culture and transfection”. The assays were performed at DIV3, 11, 18, and 21. On the day of assay, the XF assay medium was supplemented with 25 mM glucose, 0.5 mM pyruvate, and 2 mM glutamine, and the pH adjusted to 7.4. After assay performance, cells were lysed with PB1x buffer, and protein concentration was measured, and the obtained values were used for data normalization. Data analysis was performed using the Seahorse XFe Wave software and the Seahorse XF Cell Energy Phenotype Test Report Generator. The baseline OCR/ECAR ratio was higher than 4 only in DIV3, which means that the stressed ECAR parameter, in this case, may include both glycolysis and mitochondrial activity.

### Immunofluorescence staining

The animals were deeply anesthetized and transcardially perfused using saline, followed by 4% PFA. The brains were removed, post-fixed in 4% PFA for 48 h, and cryopreserved with graded sucrose (10-20-30%) over 72 h. The 20 or 30-µm parasagittal mouse brain sections were cut using a cryostat at −20°C and collected on SuperFrost Plus slides (Thermo Fisher Scientific, Waltham, MA, USA). The sections were processed immediately. The HIER procedure was applied by incubation of the sections in citrate buffer (pH 9.0) for 30 min at 60°C. The sections were blocked via incubation in 4% normal goat serum in TBS for 1h. For immunofluorescent staining, the sections were incubated overnight at 4°C with the primary rabbit anti-ataxin-3 (1:200; ProteinTech, Rosemont, IL, USA), mouse SMI-32 against non-phosphorylated heavy and medium neurofilament subunits (1:1000) and mouse SMI-312 against phosphorylated heavy and medium neurofilaments (1:1000) (Biolegends, San Diego, CA, USA), and subsequently with the anti-mouse antibody labeled by AlexaFluor488 or Dylight594 (1:400; Jackson ImmunoResearch; Suffolk, UK). The sections were end-stained with Hoechst 33342 (Sigma) nuclear stain at 1:1000 and embedded in Fluoroshield (Sigma) mounting medium.

### Image acquisition and quantification

Confocal images were acquired using 2 microscope systems. The first system was Opera LX (PerkinElmer) using 40 × water objective, 200 ms exposure time, and 50-70% laser power. For each picture, 25-50 confocal sections were collected, from which 10 sections were dissected for further analysis. The second system was TCS SP5 II (Leica Microsystems; Poland), using oil immersion 63 x objective with a sequential acquisition setting. For fluorescent quantification, images were acquired using the same settings at a resolution of 1024 x 1024 pixels and 100 Hz. Confocal sections (8-10) were acquired to a total Z-stack thickness of 0.13 μm. For each condition, we performed 3 independent cultures; and 5 independent transfections from which 46 pictures of Ki91 and 33 pictures of WT were collected and analyzed. For analyzing the state of mrFp-Rab7+ vesicles in fixed neurons, we selected axons, which were distinguished from dendrites based on known morphological characteristics: greater length and sparse branching (Banker and Cowan, 1979). For immunostaining of mouse brains, we used 4 slices, and randomly acquired from each slice, ≥ 5 fields were. Offline analysis of the image Z-stack was performed using the open-source image-processing package FIJI/Image J. Morphometric measurements were performed using FIJI/Image J. Measured data were imported into Excel software for analysis. The thresholds in all images were set to similar levels. Data were obtained from at least three independent experiments, and the number of cells or imaging sections used for quantification is indicated in the figures. All statistical analyses were performed using the Student’s t-test and are presented as mean ± SEM.

### Statistics

The data regarding behavioral experiments were subjected to a two-way ANOVA, followed by Bonferroni post-tests. P-values of less than 0.05 were considered significant. Identification of proteins on raw proteomic data was performed by the Andromeda search engine in Mascot using the following inclusion criteria: 1. At least two different peptides per protein were identified per sample, and a total peptide score reached the significance threshold FDR=0.01. Identified proteins matching the inclusion criteria were subjected to further statistical analysis with a two-sample t-test, and dysregulation of protein level reaching p-value < 0.05 was considered as significant. Kruskal-Wallis test was used to perform a score assessment in the 0-5 scale of mild, moderate, and severe phenotype in 18-month-old animals (p-value < 0.05).

### Availability of data and materials

The complete and processed datasets are available along with the manuscript as supplementary material, while the row data will be available from the corresponding author on request.

## Acknowledgments

Proteomic mass spectrometry analyses were performed in the Laboratory of Mass Spectrometry (European Centre for Bioinformatics and Genomics; ECBiG; IBCH, PAS, Poznań, Poland). Confocal imaging was performed using the facilities in the Laboratory of Subcellular Structures Analysis and Laboratory of Molecular Assays (IBCh, PAS, Poland). We thank Adam Plewinski for his help in maintaining the colonies of mice. We thank Joanna Mieloch for help in maintaining the animals. We thank Magdalena Surdyka and Łukasz Przybył for collecting tissues, which were sent for MRI.

## Abbreviations

SCA3: Spinocerebellar ataxia type 3

MJD: Machado-Joseph disease

polyQ: polyglutamine

MS: mass spectrometry

PCA: principal component analysis

ANOVA: analysis of variance

FC: fold change

WT: wildtype

mo: month-old

CPDB: ConsensusPath Database

BP: biological process

MF: molecular function

CC: cellular compartment

mRFP: monomeric red fluorescent protein

OCR: oxygen consumption rate

ECAR: extracellular acidification rate

TCA: tricarboxylic acid cycle

OXPHOS: oxidative phosphorylation

CNS: central nervous system

MRI: magnetic resonance imaging

mRNP: messenger ribonucleoprotein

DIV: days in vitro

NF: neurofilament

## Author contributions statement

MF conceived, designed, supervised all experiments, and analyzed the data. KW and ŁM performed all proteomic experiments. KW designed and performed all experiments except of MRI. EB, JB, JF performed the MRI experiment and analyzed the data. MF and KW wrote the paper. MF was responsible for the research concept and obtaining funding.

## Supplementary material

**Supplementary Table 1.** MS/MS proteomic data for each tested age (4, 10, 12, 14 mo) in the parallel approach in the cerebellum (excel file)

**Supplementary Table 2.** MS/MS proteomic data for each tested age (4, 10, 12, 14 mo) in the parallel approach in the cerebral cortex (excel file)

**Supplementary Table 3.** MS/MS proteomic data for 18-month-old Ki91 cerebellum and cortex in the correlative approach (excel file)

**Supplementary Table 4.** MS/MS proteomic data for the of protein levels in somatodendritic and axon compartment in cerebellar and cortical samples (excel file)

**Supplementary Table 5** Proteins depleted in Ki91 cerebellar axons Supplementary Table 6 Proteins enriched in Ki91 cerebellar axons Supplementary Table 7 Proteins depleted in Ki91 cortical axons Supplementary Table 8 Proteins enriched in Ki91 cortical axons

---

**Supplementary Figure 1.**
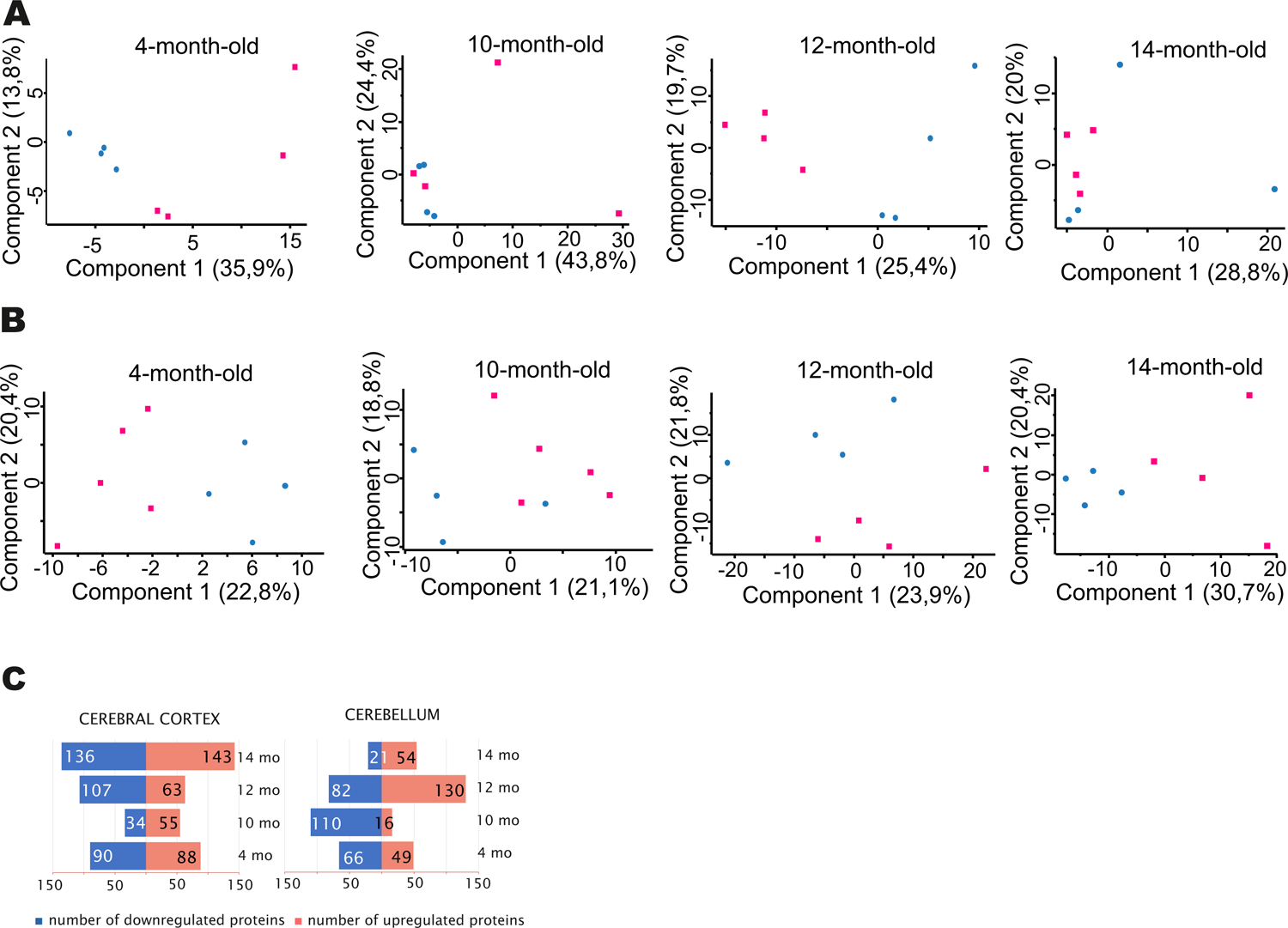
Principal Component Analysis for proteomic data in the “parallel” approach of the cerebellum and cerebral cortex samples. Distinct clustering of dysregulated genes of proteomic data of the cerebellum (A) and cerebral cortex (B) of K91 and C57BL/6 mice (n=4-11 per genotype) is presented in the form of PCA graphs. The principal component analysis was performed in Perseus software ver. 1.6.1.3. Pink color denotes Ki91 brain samples, blue color C57BL/6 brain samples. A total number of 1058 dysregulated proteins were identified in the cerebral cortex and 830 in the cerebellum (p<0.05; two-sample t-test) (C). There was no clear trend neither towards downregulation or upregulation.

**Supplementary Figure 2.**
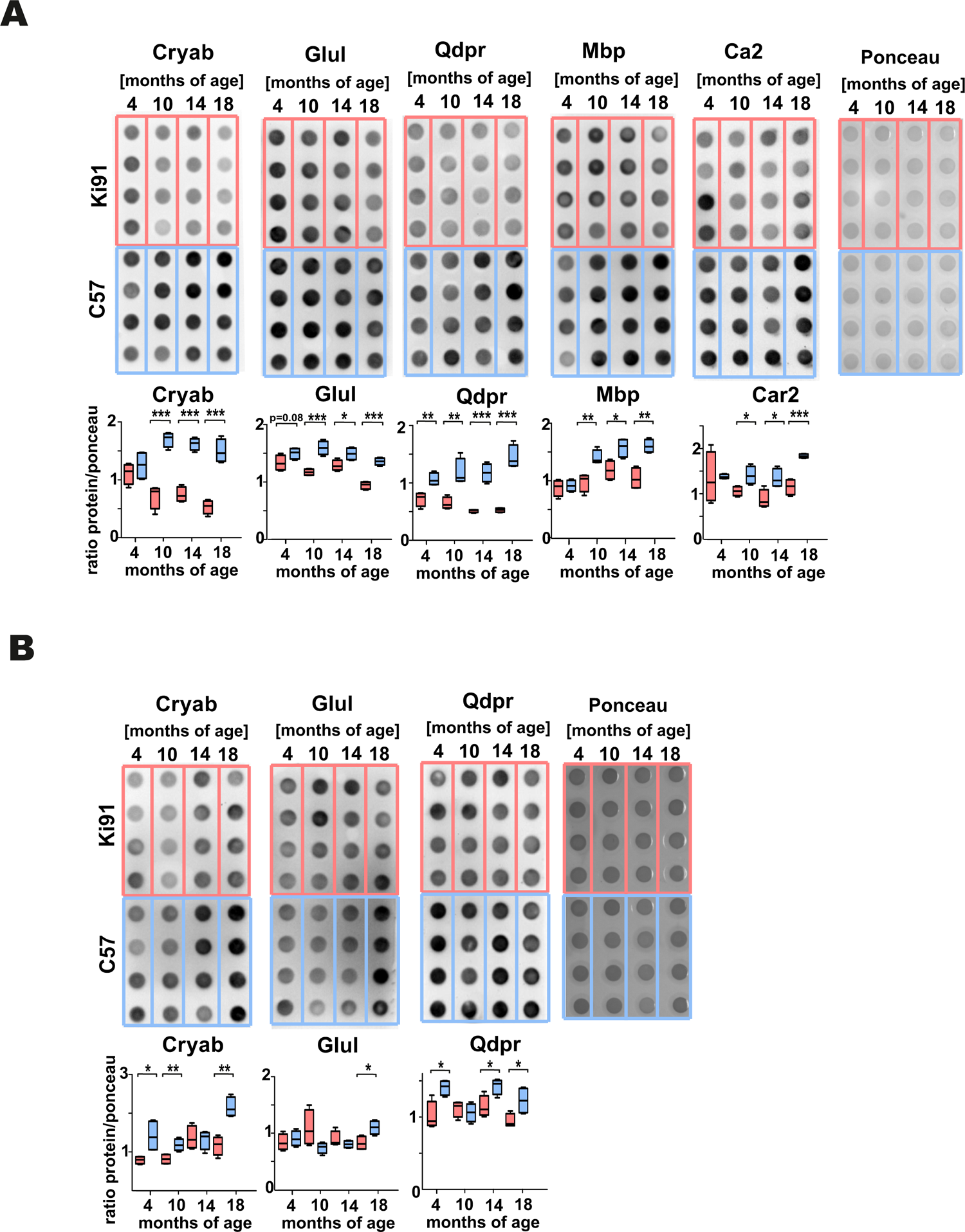
Validation of dysregulated proteins identified in the LC-MS/MS proteomic “parallel” approach with vacuum dot blot assay in the Ki91 SCA3/MJD brain tissues. Vacuum dot blot analysis confirmed decreased levels of Cryab (p < 0.001; two-sample t-test), Glul (p <0.05; two-sample t-test), Mbp (p<0.05; two-sample t-test) and Car2 (p<0.05; two-sample t-test) in the cerebral cortex of 10, 14, and 18-month-old Ki91 mice and Qdpr (p < 0.01; two-sample t-test) in the 4, 10, 14, and 18-month-old Ki91 mice (A). In the cerebellum of Ki91 animals, decreased levels of Cryab (p <0.05; two-sample t-test) and Qdpr (p < 0.01; two-sample t-test) was shown in 4, 10, and 18-month-old Ki91 mice and decreased levels of Glul in 18-month-old mice (F). Ponceau was used as a loading control. N=4 per genotype, error bars: SEM. All experiments were performed in 3 technical replicates.

**Supplementary Figure 3.**
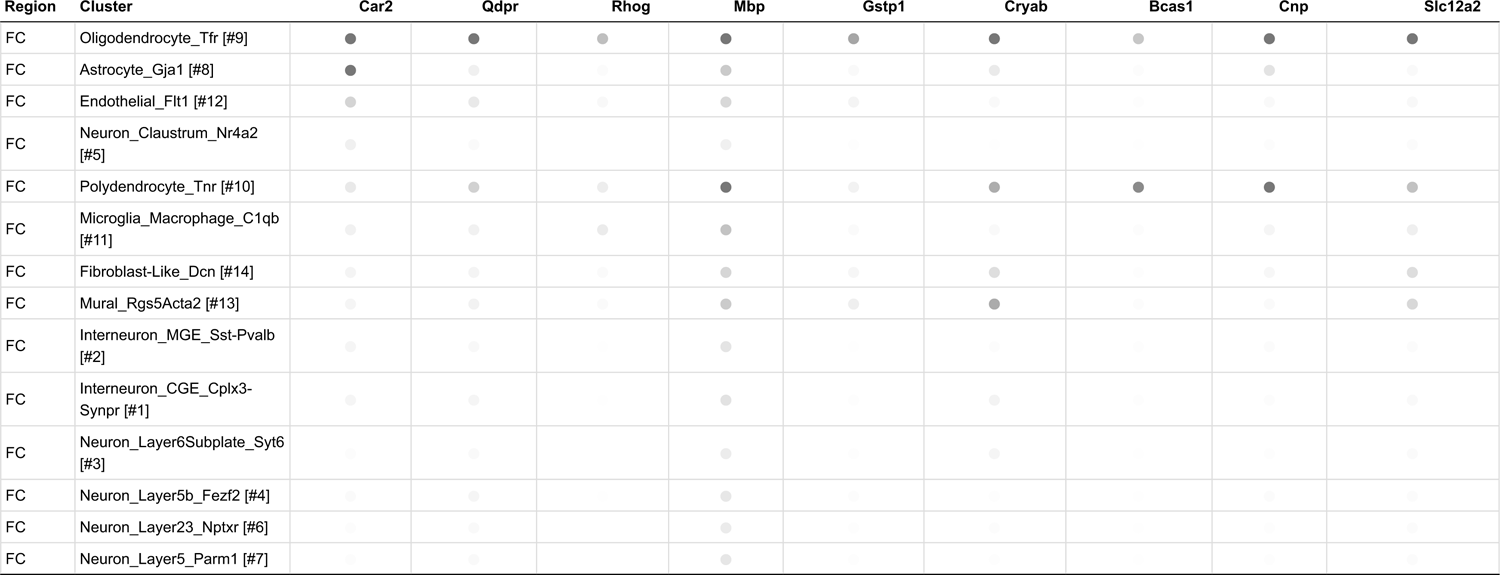
Cell types affected by dysregulation of proteins in the cerebral cortex of Ki91 SCA3/MJD mice The identification of cell types was performed with Dropviz (http://dropviz.org) for dysregulated proteins. In the cerebellar cortex, we identified a group of dysregulated proteins with high expression in oligodendrocytes and polydendrocytes. Among these proteins were Ca2, Odpr, Rhog, Mbp, Gstp1, Cryab, Bcas1, Cnp, and Slc12a2. The protein was considered as a cellular marker if the ratio of its level to the level of expression in other cell types was 3 or more. Darker dots represent higher gene expression (transcripts per 100 000 in the cluster (TPT).

**Supplementary Figure 4.**
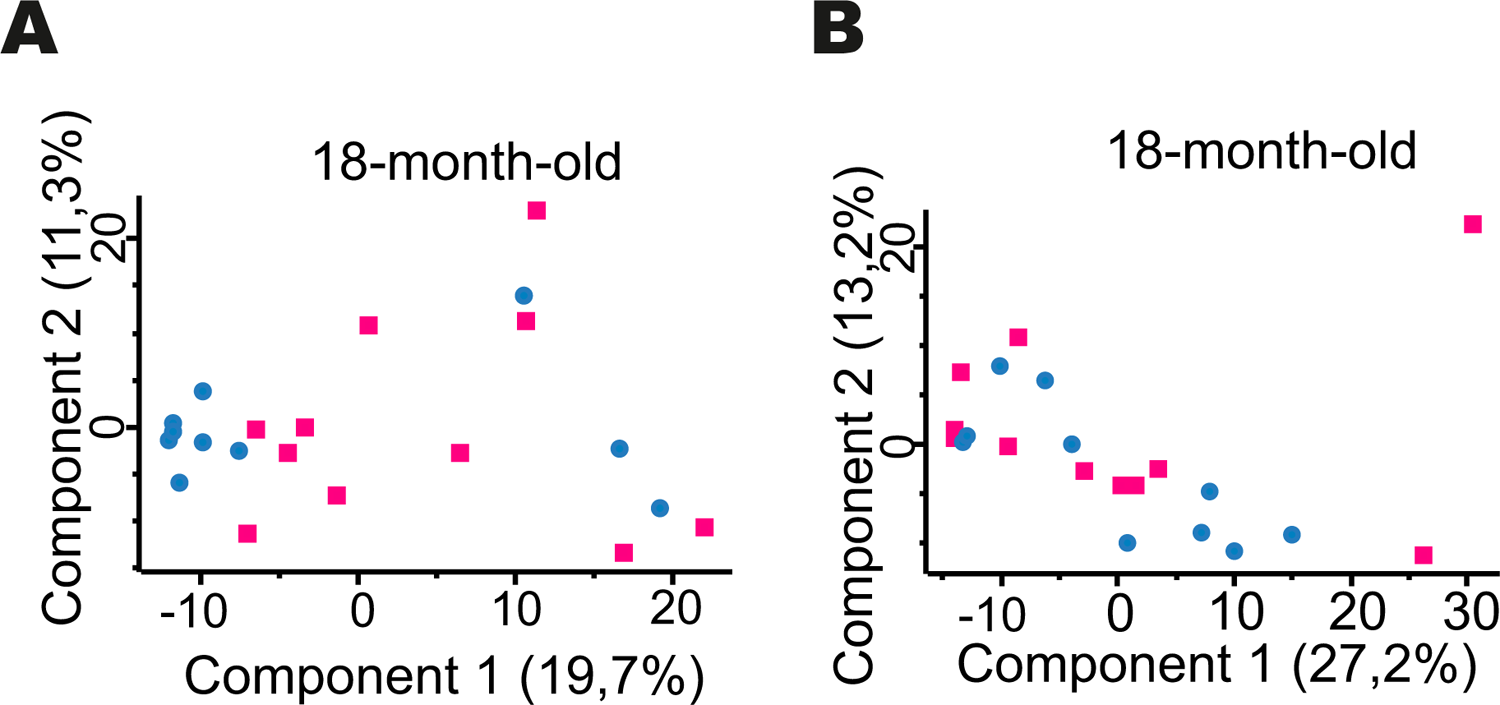
Principal Component Analysis for proteomic data in the correlative approach of the cerebellum and cerebral cortex samples A total number of 2555 proteins were identified in the cerebral cortex and cerebellum of 18-month-old mice (FDR <0.01). Distinct clustering of dysregulated genes of proteomic data of the cerebellum (A) and cerebral cortex (B) of K91 and C57BL/6 mice (n = 10 WT C57BL/6 and n=11 Ki91 mut/mut) is presented in the form of PCA graphs. The principal component analysis was performed in Perseus software ver. 1.6.1.3. Pink color denotes Ki91 brain samples, blue color C57BL/6 brain samples.

**Supplementary Figure 5.**
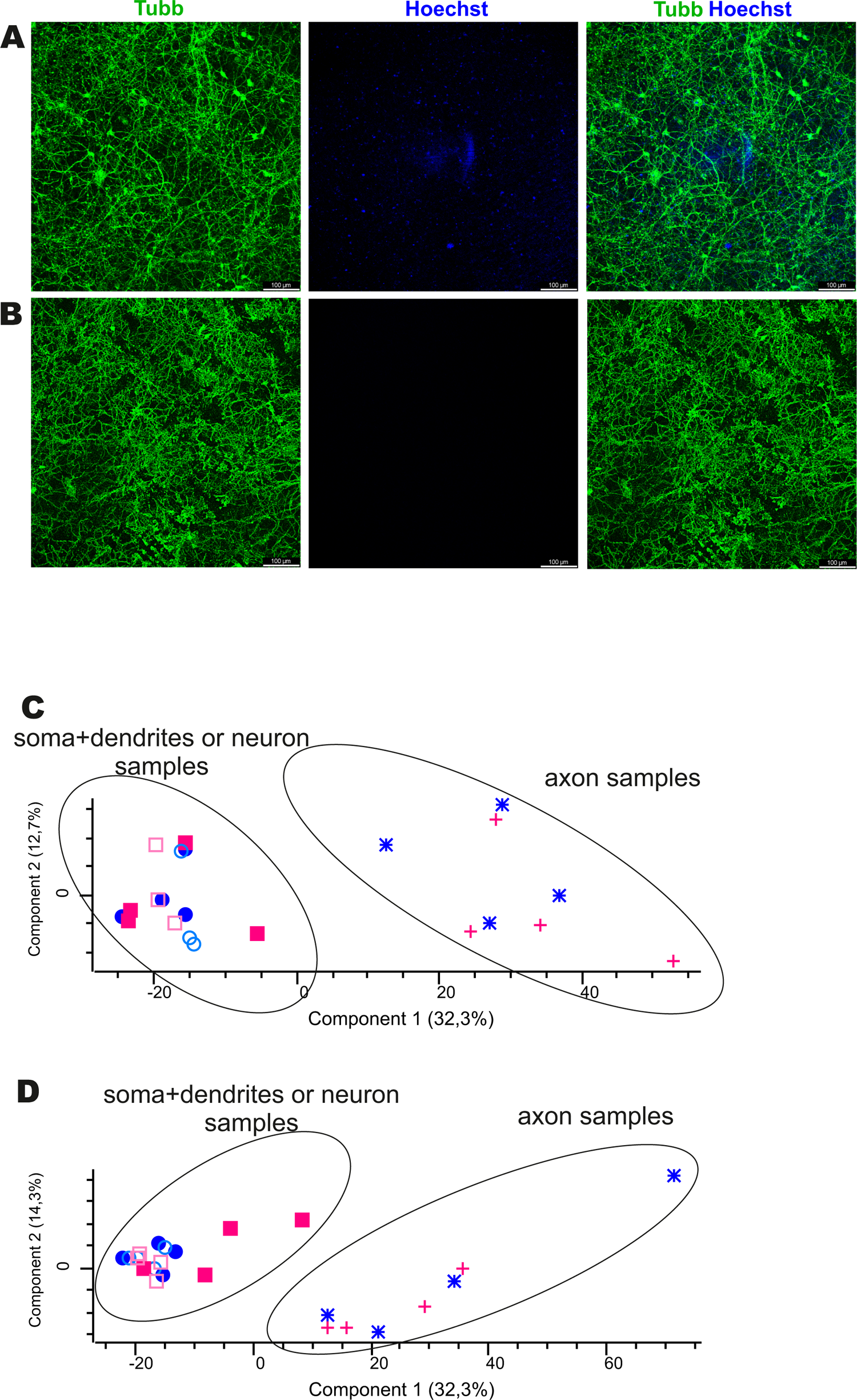
Quality control of proteomic analysis of axonal transport Clear separation of an axonal fraction from the somatodendritic fraction is demonstrated by immunostaining of culture filters isolated from the Boyden chamber before (A) and after scraping off the cell bodies (B), with the antibody for B-Tubulin (green) and nuclear dye Hoechst (blue). In addition, PCA graphs also show a clear separation between axonal and somatodendritic samples from cerebellar neurons (C) and cortical neurons (D).

**Supplementary Figure 6.**
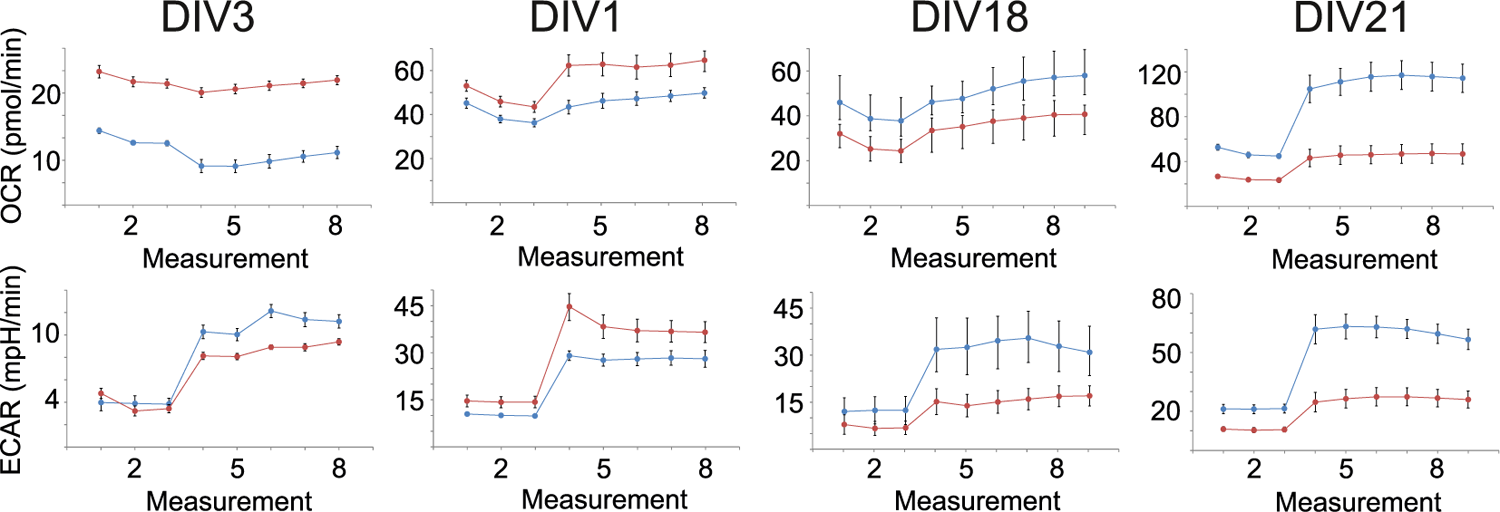
Seahorse XFp Cell Energy Phenotype profiling in cerebellar neurons The rate of mitochondrial respiration (OCR, oxygen consumption rate) and glycolysis (ECAR, extracellular acidification rate) were measured under baseline (3 measurements) and stressed conditions (5 measurements), which were caused by the injection of 1 μM of oligomycin and 1 μM of FCCP). The profiles of OCR (upper panel) and ECAR (lower panel) during testing are presented on the graphs.

