## Supplementary Table 5 and 6 for "Broad influence of mutant ataxin-3 on the proteome of the adult brain, young neurons, and axons reveals central molecular processes and biomarkers in SCA3/MJD using knock-in mouse model": table 5 and 6.docx

Table 5. Proteins depleted in Ki91 cerebellar axons

| Category | GO term | GO ID | | | q value (GO term) | | | protein | p value | Ratio axon to soma |
| --- | --- | --- | --- | --- | --- | --- | --- | --- | --- | --- |
| Cytoskeleton | | | |  |  | | |  |  |  |
| actin cytoskeleton (CC level 5) | | | | GO:0015629 | 3.37e-05 | | | Flna | 0.023 | 0.28 |
| actin cytoskeleton (CC level 5) | | | | GO:0015629 | 3.37e-05 | | | Myl6 | 0.001 | 0.66 |
| actin cytoskeleton (CC level 5) | | | | GO:0015629 | 3.37e-05 | | | Sptan1 | 0.037 | 0.63 |
| actin cytoskeleton (CC level 5) | | | | GO:0015629 | 3.37e-05 | | | Sptbn1 | 0.031 | 0.57 |
| actin cytoskeleton (CC level 5) | | | | GO:0015629 | 3.37e-05 | | | Wasf1 | 0.037 | 0.50 |
| actin cytoskeleton (CC level 5) | | | | GO:0015629 | 3.37e-05 | | | Coro1b | 0.012 | 0.51 |
| actin cytoskeleton (CC level 5) | | | | GO:0015629 | 3.37e-05 | | | Capn2 | 0.035 | 0.06 |
| cell cortex (CC level 5) | | | | GO:0005938 | 1.69e-06 | | | Ctnnb1 | 0.010 | 0.44 |
| cytoskeleton (CC level 4) | | | | GO:0005856 | 0.00246 | | | Ank2 | 0.025 | 0.49 |
| cytoskeleton (CC level 4) | | | | GO:0005856 | 0.00246 | | | Mapk1 | 0.024 | 0.52 |
| cytoskeleton (CC level 4) | | | | GO:0005856 | 0.00246 | | | Tpr | 0.016 | 0.34 |
| Filament-forming cytoskeletal GTPase | | | | UniProtKB | - | | | Septin3 | 0.014 | 0.62 |
| alpha-tubulin binding | | | | GO:0043014 | - | | | Sncb | 0.003 | 0.65 |
| Transport | | | |  |  | | |  |  |  |
| vesicle-mediated transport (BP level 4) | | | | GO:0016192 | 0.0008 | | | Actr1a | 0.006 | 0.74 |
| vesicle-mediated transport (BP level 4) | | | | GO:0016192 | 0.0008 | | | Arf6 | 0.012 | 0.72 |
| vesicle-mediated transport (BP level 4) | | | | GO:0016192 | 0.0008 | | | Nsf | 0.029 | 0.69 |
| vesicle-mediated transport (BP level 4) | | | | GO:0016192 | 0.0008 | | | Rhob | 0.041 | 0.55 |
| Vesicles | | | |  |  | | |  |  |  |
| SNARE complex (CC level 3) | | | | GO:0031201 | 0.0104 | | | Napa | 0.029 | 0.49 |
| SNARE complex (CC level 3) | | | | GO:0031201 | 0.0104 | | | Vti1b | 0.044 | 0.30 |
| Vesicle (CC level 3) | | | | GO:0031982 | 0.0021 | | | Sv2b | 0.015 | 0.57 |
| Mitochondria | | | |  |  | | |  |  |  |
| mitochondrion (CC level 4) | | | | GO:0005739 | 6.73e-05 | | | Bnip3 | 0.020 | 0.67 |
| mitochondrion (CC level 4) | | | | GO:0005739 | 6.73e-05 | | | Fech | 0.030 | 0.57 |
| mitochondrion (CC level 4) | | | | GO:0005739 | 6.73e-05 | | | Ak4 | 0.021 | 0.32 |
| mitochondrion (CC level 4) | | | | GO:0005739 | 6.73e-05 | | | Pccb | 0.029 | 0.68 |
| mitochondrion (CC level 4) | | | | GO:0005739 | 6.73e-05 | | | Gfm1 | 0.042 | 0.40 |
| mitochondrion (CC level 4) | | | | GO:0005739 | 6.73e-05 | | | Gdap1 | 0.029 | 0.37 |
| mitochondrion (CC level 4) | | | | GO:0005739 | 6.73e-05 | | | Mtch1 | 0.025 | 0.43 |
| mitochondrion (CC level 4) | | | | GO:0005739 | 6.73e-05 | | | Txn2 | 0.033 | 0.41 |
| Membrane | | | |  |  | | |  |  |  |
| cell body membrane (CC level 3) | | | | GO:0044298 | 0.00453 | | | Kcnd2 | 0.00022 | 0.62 |
| cell body membrane (CC level 3) | | | | GO:0044298 | 0.00453 | | | Slc4a8 | 0.038 | 0.45 |
| synaptic membrane (CC level 2) | | | | GO:0097060 | 0.0188 | | | Cadm3 | 0.035 | 0.55 |
| Other | | | |  |  | | |  |  |  |
| endomembrane system (CC level 2) | | | | GO:0012505 | 0.0155 | | | Ei24 | 0.039 | 0.60 |
| endomembrane system (CC level 2) | | | | GO:0012505 | 0.0155 | | | Pgrmc2 | 0.025 | 0.64 |
| endomembrane system (CC level 2) | | | | GO:0012505 | 0.0155 | | | Rdh11 | 0.004 | 0.36 |
| endomembrane system (CC level 2) | | | | GO:0012505 | 0.0155 | | | Ccdc136 | 0.021 | 0.32 |
| intracellular protein transport (BP level 4) | | | | GO:0006886 | 0.00111 | | | Rpl15 | 0.046 | 0.39 |
| cell differentiation with neurite sprouting | | | | Entrez | - | | Gdap1l1 | | 0.011 | 0.34 |
| creatine kinase activity | | | | GO:0004111 | - | | Ckmt1 | | 0.009 | 0.32 |
| protein binding involved in protein folding | | | GO:0044183 | | | - | | Pfdn1 | 0.005 | 0.17 |
| pre-mRNA splicing | | | UniProtKB | | | - | | Snrpd2 | 0.001 | 0.08 |

Table 6. Proteins enriched in Ki91 cerebellar axons

| Category | GO term | GO ID | | q value (GO term) | protein | p value | Ratio axon to soma |
| --- | --- | --- | --- | --- | --- | --- | --- |
| Ribosme and RNA-binding proteins | | |  |  |  |  |  |
| cytosolic ribosome (CC level 4) | | | GO:0022626 | 2.69e-05 | Rpl22l1 | 0.048 | 4.78 |
| cytosolic ribosome (CC level 4) | | | GO:0022626 | 2.69e-05 | Rpl13 | 0.0004 | 2.00 |
| cytosolic ribosome (CC level 4) | | | GO:0022626 | 2.69e-05 | Rpl35a | 0.030 | 3.56 |
| cytosolic ribosome (CC level 4) | | | GO:0022626 | 2.69e-05 | Rps5 | 0.038 | 2.77 |
| ribosome (CC level 3) | | | GO:0005840 | 0.000102 | Rrbp1 | 0.032 | 2.94 |
| RNA binding (MF level 4) | | | GO:0003723 | 0.000141 | Snu13 | 0.023 | 2.09 |
| RNA binding (MF level 4) | | | GO:0003723 | 0.000141 | Pcbp3 | 0.012 | 3.02 |
| RNA binding (MF level 4) | | | GO:0003723 | 0.000141 | Zfr | 0.030 | 3.60 |
| regulation of mRNA stability (BP level 4) | | | GO:0043488 | 0.00105 | Npm1 | 0.034 | 2.28 |
| regulation of mRNA stability (BP level 4) | | | GO:0043488 | 0.00105 | Cirbp | 0.028 | 4.92 |
| regulation of mRNA stability (BP level 4) | | | GO:0043488 | 0.00105 | Fxr2 | 0.029 | 2.80 |
| regulation of mRNA stability (BP level 4) | | | GO:0043488 | 0.00105 | Hspa8 | 0.021 | 1.39 |
| Proteasome | | |  |  |  |  |  |
| proteasome complex (CC level 3) | | | GO:0000502 | 0.0195 | Psmd14 | 0.015 | 10.75 |
| proteasome complex (CC level 3) | | | GO:0000502 | 0.0195 | Psma5 | 0.035 | 1.60 |
| Cytoskeleton | | |  |  |  |  |  |
| Filament-forming cytoskeletal GTPase | | | UniProtKB | - | Septin4 | 0.005 | 4.06 |
| Actin-binding protein regulating the actin cytoskeleton | | | Entrez | - | Cotl1 | 0.041 | 2.06 |
| Other | | |  |  |  |  |  |
| extracellular vesicle (CC level 3) | | | GO:1903561 | 0.00361 | Igsf8 | 0.038 | 1.64 |
| extracellular vesicle (CC level 3) | | | GO:1903561 | 0.00361 | Basp1 | 0.031 | 2.37 |
| extracellular vesicle (CC level 3) | | | GO:1903561 | 0.00361 | Serinc5 | 0.008 | 1.55 |
| chaperone cofactor-dependent protein refolding (BP level 4) | | | GO:0051085 | 0.00924 | Ero1a | 0.038 | 4.06 |
| cellular response to topologically incorrect protein (BP level 4) | | | GO:0035967 | 0.0133 | Sdf2 | 0.019 | 1.56 |
| intracellular membrane-bounded organelle (CC level 3) | | | GO:0043231 | 0.0119 | Naxd | 0.015 | 1.91 |
| intracellular membrane-bounded organelle (CC level 3) | | | GO:0043231 | 0.0119 | Lamp1 | 0.006 | 11.08 |
| intracellular membrane-bounded organelle (CC level 3) | | | GO:0043231 | 0.0119 | Lrpap1 | 0.040 | 5.16 |
| intracellular membrane-bounded organelle (CC level 3) | | | GO:0043231 | 0.0119 | Mif | 0.014 | 1.98 |
| intracellular membrane-bounded organelle (CC level 3) | | | GO:0043231 | 0.0119 | Sfxn5 | 0.015 | 1.98 |
| intracellular membrane-bounded organelle (CC level 3) | | | GO:0043231 | 0.0119 | Mecr | 0.027 | 1.74 |
| intracellular membrane-bounded organelle (CC level 3) | | | GO:0043231 | 0.0119 | Eps15l1 | 0.016 | 3.89 |
| intracellular membrane-bounded organelle (CC level 3) | | | GO:0043231 | 0.0119 | Stx12 | 0.019 | 1.44 |
| intracellular membrane-bounded organelle (CC level 3) | | | GO:0043231 | 0.0119 | Cntnap2 | 0.007 | 4.65 |
| intracellular membrane-bounded organelle (CC level 3) | | | GO:0043231 | 0.0119 | Cyp2s1 | 0.038 | 1.64 |
| Calcium ion binding | | | GO:0005509 | - | Psph | 0.030 | 3.64 |
| Cysteamine dioxygenase activity | | | GO:0017172 | - | Ado | 0.0006 | 1.92 |
| Nerve growth | | | UniProtKB | - | Gap43 | 0.012 | 1.97 |
