## Supplementary Table 7 and 8 for "Broad influence of mutant ataxin-3 on the proteome of the adult brain, young neurons, and axons reveals central molecular processes and biomarkers in SCA3/MJD using knock-in mouse model": Table 7 and 8.docx

Table 7. Proteins depleted in Ki91 cortical axons

| Category | GO term | GO ID | q value (GO term) | protein | p value | Ratio axon to soma |
| --- | --- | --- | --- | --- | --- | --- |
| Cytoskeleton | |  |  |  |  |  |
| actin filament bundle organization (BP level 5) | | GO:0061572 | 0.123 | Pfn1 | 0.028 | 0.57 |
| actin filament bundle organization (BP level 5) | | GO:0061572 | 0.123 | Shtn1 | 0.022 | 0.09 |
| structural constituent of cytoskeleton | | GO:0005200 | - | Nefm | 0.046 | 0.54 |
| Golgi apparatus | |  |  |  |  |  |
| Golgi membrane (CC level 4) | | GO:0000139 | 0.0758 | Tmco1 | 0.021 | 0.26 |
| Golgi membrane (CC level 4) | | GO:0000139 | 0.0758 | Trappc3 | 0.017 | 0.50 |
| Golgi membrane (CC level 4) | | GO:0000139 | 0.0758 | Gorasp2 | 0.048 | 0.32 |
| Golgi membrane (CC level 4) | | GO:0000139 | 0.0758 | Scamp5 | 0.034 | 0.78 |
| Mitochondria | |  |  |  |  |  |
| mitochondrial membrane organization (BP level 4) | | GO:0007006 | 0.121 | Mfn2 | 0.008 | 0.68 |
| mitochondrial membrane organization (BP level 4) | | GO:0007006 | 0.121 | Timm13 | 0.009 | 0.44 |
| electron transfer activity and NAD binding | | GO:0009055 | - | Me1 | 0.047 | 0.63 |
| Protein localization | |  |  |  |  |  |
| protein localization to membrane (BP level 5) | | GO:0072657 | 0.117 | Dpp6 | 0.046 | 0.71 |
| protein localization to membrane (BP level 5) | | GO:0072657 | 0.117 | Rps28 | 0.047 | 0.53 |
| protein localization to membrane (BP level 5) | | GO:0072657 | 0.117 | Timm13 | 0.009 | 0.44 |
| cellular protein localization (BP level 4) | | GO:0034613 | 0.121 | Nup62 | 0.036 | 0.20 |
| Other | |  |  |  |  |  |
| bounding membrane of organelle (CC level 3) | | GO:0012505 | 0.0155 | Tmem35a | 0.018 | 0.54 |
| cadherin binding (MF level 4) | | GO:0012505 | 0.0192 | Prdx1 | 0.042 | 0.75 |
| calcium ion binding | | GO:0005509 | - | Capns1 | 0.046 | 0.60 |
| endonuclease activity | | GO:0004519 | - | Exog | 0.013 | 0.36 |
| ubiquitination | | UniProtKB | - | Sugt1 | 0.0009 | 0.16 |
| protein homodimerization activity | | GO:0046983 | - | Cbs | 0.005 | 0.14 |
| - | | - | - | Ccdc58 | 0.041 | 0.45 |

Table 8. Proteins enriched in Ki91 cortical axons

| Category | GO term | GO ID | | q value (GO term) | protein | p value | Ratio axon to soma |
| --- | --- | --- | --- | --- | --- | --- | --- |
| Translation and splicing | | |  |  |  |  |  |
| translation preinitiation complex (CC level 3) | | | GO:0070993 | 6.16e-05 | Eif3e | 0.025 | 16.02 |
| translation preinitiation complex (CC level 3) | | | GO:0070993 | 6.16e-05 | Eif3g | 0.037 | 2.14 |
| translation preinitiation complex (CC level 3) | | | GO:0070993 | 6.16e-05 | Eif3h | 0.046 | 2.10 |
| ribosome (CC level 3) | | | GO:0005840 | 0.000237 | Rpl8 | 0.043 | 1.79 |
| ribosome (CC level 3) | | | GO:0005840 | 0.000237 | Rpl19 | 0.048 | 2.92 |
| ribosome (CC level 3) | | | GO:0005840 | 0.000237 | Rpl28 | 0.029 | 3.42 |
| ribosome (CC level 3) | | | GO:0005840 | 0.000237 | Mrpl48 | 0.034 | 2.32 |
| Binds RNA | | | UniProtKB | - | Pa2g4 | 0.023 | 1.73 |
| U2-type spliceosomal complex (CC level 4) | | | GO:0005684 | 0.00116 | Snrpd2 | 0.022 | 3.32 |
| U2-type spliceosomal complex (CC level 4) | | | GO:0005684 | 0.00116 | Luc7l3 | 0.040 | 2.83 |
| U2-type spliceosomal complex (CC level 4) | | | GO:0005684 | 0.00116 | Smu1 | 0.002 | 2.64 |
| Protein transport and localization | | |  |  |  |  |  |
| intracellular protein transport (BP level 4) | | | GO:0006886 | 0.00685 | Sel1l | 0.027 | 3.35 |
| intracellular protein transport (BP level 4) | | | GO:0006886 | 0.00685 | Surf4 | 0.031 | 5.44 |
| intracellular protein transport (BP level 4) | | | GO:0006886 | 0.00685 | Tmed4 | 0.001 | 6.21 |
| establishment of protein localization (BP level 4) | | | GO:0045184 | 0.0113 | Pdcd6ip | 0.035 | 4.63 |
| establishment of protein localization (BP level 4) | | | GO:0045184 | 0.0113 | Cct2 | 0.044 | 1.82 |
| Other | | |  |  |  |  |  |
| vesicle (CC level 3) | | | GO:0031982 | 0.00228 | Lgals1 | 0.031 | 3.90 |
| vesicle (CC level 3) | | | GO:0031982 | 0.00228 | Tmem33 | 9.80e-05 | 4.32 |
| lytic vacuole (CC level 5) | | | GO:1903561 | 0.00409 | Tpp1 | 0.043 | 2.30 |
| lytic vacuole (CC level 5) | | | GO:0000323 | 0.00409 | Atp8a1 | 0.040 | 2.17 |
| carbohydrate transmembrane transport  (BP level 5) | | | GO:0034219 | 0.0456 | Pea15 | 0.045 | 2.59 |
| intracellular membrane-bounded organelle (CC level 3) | | | GO:0043231 | 0.0228 | Osbpl8 | 0.037 | 9.88 |
| intracellular membrane-bounded organelle (CC level 3) | | | GO:0043231 | 0.0228 | Ndufb8 | 0.034 | 4.95 |
| intracellular membrane-bounded organelle (CC level 3) | | | GO:0043231 | 0.0228 | Tcea1 | 0.019 | 2.22 |
| intracellular membrane-bounded organelle (CC level 3) | | | GO:0043231 | 0.0228 | Ncln | 0.037 | 2.06 |
| G protein-coupled GABA receptor activity | | | GO:0004965 | - | Gabbr2 | 0.017 | 2.95 |
| - | | | - | - | Fsd1l | 0.033 | 2.05 |
